# A generic framework for hierarchical *de novo* protein design

**DOI:** 10.1101/2022.04.07.487481

**Authors:** Zander Harteveld, Jaume Bonet, Stéphane Rosset, Che Yang, Fabian Sesterhenn, Bruno E. Correia

**Affiliations:** Institute of Bioengineering, École Polytechnique Fédérale de Lausanne, Lausanne, CH-1015, Switzerland; Swiss Institute of Bioinformatics (SIB), Lausanne CH-1015, Switzerland

## Abstract

*De novo* protein design enables the exploration of novel sequences and structures absent from the natural protein universe. *De novo* design also stands as a stringent test to our understanding of the underlying physical principles of protein folding and may lead to the development of proteins with unmatched functional characteristics. The first fundamental challenge of *de novo* design is to devise “designable” structural templates leading to sequences that will adopt the predicted fold. Here, we built on the TopoBuilder *de novo* design method to automatically assemble structural templates with native-like features starting from string descriptors that capture the overall topology of proteins. Our framework eliminates the dependency of hand-crafted and fold-specific rules through an iterative, data-driven approach that extracts geometrical parameters from structural tertiary motifs. We evaluated the TopoBuilder framework by designing sequences for a set of five protein folds and experimental characterization revealed that several sequences were folded and stable in solution. The TopoBuilder *de novo* design framework will be broadly useful to guide the generation of artificial proteins with customized geometries, enabling the exploration of the protein universe.

## Introduction

Evolution has only explored a small subset of all possible amino acid (AA) sequences and structures [1]. The space of viable protein sequences e.g., sequences that have a global free energy minimum representing a well-folded native state, is small. Such a notion has been supported by several experimental studies showing that most random AA sequences have a rough energy landscape with many local minima representing aggregated or misfolded states [2]–[5].

*De novo* design strategies stand as an essential tool to aid the exploration of the sequence space and thereby enabling the creation of new protein structures and functions. Classical *de novo* protein design generally entails two iterative steps: first, target folds are modeled (backbone generation); second, AA sequences that stabilize the lowest free energy state of the target backbone conformation are searched (sequence design). Despite multiple successes [6]–[9], *de novo* design remains a challenging problem for protein designers given that it stands as a stringent test to our understanding of the principles that govern protein structures.

Successful structure-based *de novo* design largely relies on the crafting of “designable” protein backbones, meaning physically realistic and strainless backbones that are compatible with sequences that will yield a protein fold with a well-defined energy minimum [10]–[15]. The designability of a protein backbone is generally proxied by the number of sequences a backbone can support [10], [16], [17]. For example, some natural protein structures can accommodate more sequences than the average and are thought to be more robust against random mutations, and therefore thermodynamically more stable which favors evolutionary stability [16]. Generating designable backbones is important as one would like to *a priori* limit the sampling space to engineer only reasonable shapes with inherent structure-to-sequence compatibility and discard presumptive non-viable structures. Many *de novo* design approaches are likely to fail due to the lack of designability of the starting structural templates, requiring multiple iterative rounds of human-guided and experimental optimizations [4], [5].

Quantifying the designability of protein backbones is difficult [18], [19]. Even recent energy functions fail to reliably capture global designability aspects of protein backbones but excel in assessing high-resolution details such as van der Waals forces, steric repulsion, electrostatic interactions, and hydrogen bonds [20]. To facilitate the *de novo* design process at early stages, it would be necessary to have low-resolution energy functions that could accurately capture the physicochemical determinants of realistic structures at the backbone level [21].

There has been considerable progress in developing parametric functions and general principles for describing ideal and less symmetric protein structures [17], [22], [23]. Often secondary structure elements (SSEs) are connected to create tertiary structural topologies by packing α-helices on paired β-strands through the control of the loop length and ABEGO residue torsion structure. This approach made it possible to design a set of ideal protein structures, including TIM-barrels [24], β-barrels [25], [26], Jelly-rolls [27], and immunoglobulin-like domains [28]. Nonetheless, parametric definitions are often specifically framed for distinct protein classes or architecture types and cannot be generalized to other architectural configurations i.e. the Crick coiled-coil generating equations [29] or descriptive parametric models of β-barrels [30], [31].

In this work, we enhanced the capabilities of the TopoBuilder framework by introducing a data-driven correction module to generate native-like backbones from a simplified description of a protein topology that we term “Sketch” (Fig. 1A and Supp. Fig. S1). This module is applicable to any protein topology that can be described by arranging ideal SSEs in layers [32], [33]. The correction module generates parametric refinements that geometrically optimize the SSEs of protein backbones towards native-like configurations, rendering them more designable. The set of corrections includes translational and rotational parameters jointly capturing key geometric features such as distances and angles of native tertiary motifs. To further aid the topology assembly step, we use structural fragments from naturally occurring loops to connect two subsequent SSEs. We evaluated the general framework by *de novo* designing five different folds and found that even a minimal set of corrections to the protein backbones is sufficient to improve sequence sampling and achieve a better sequence-to-structure compatibility and sequence quality overall according to a variety of computational metrics. Finally, we experimentally characterized 54 designs and obtained multiple sequences that were folded and stable in solution, including topologies that have been particularly difficult for computational design such as all-β structures.

**Figure 1.**
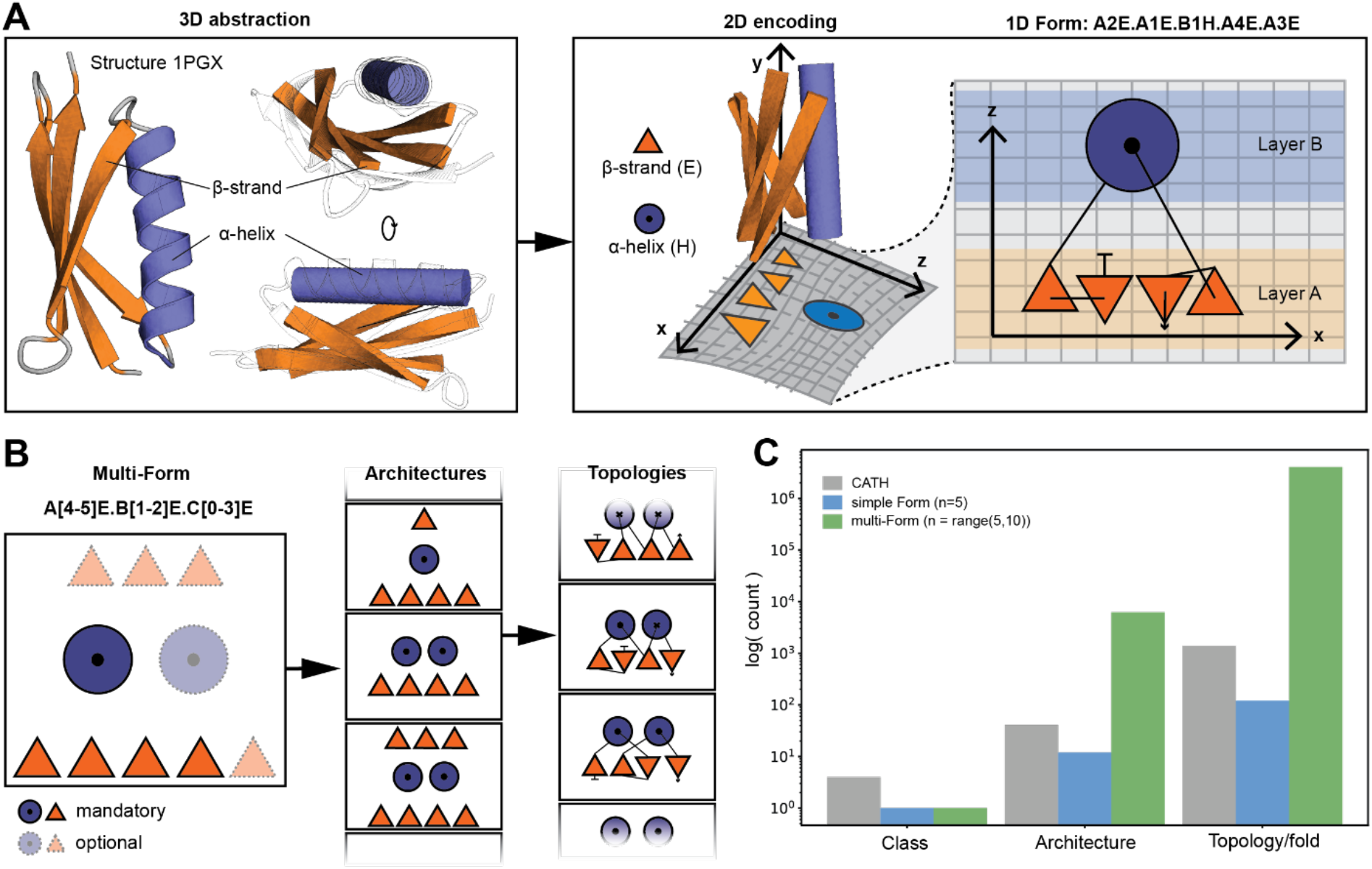
Form parametrization for protein structures. **A:** Form parametrization for a Ubiquitin-like fold (PDB 1PGX). A 3D abstraction of the structure is created that is encoded into a layered 2D lattice diagram where the sheet is assigned as layer A and the helix as layer B (layer assignment is arbitrary). The SSEs on a layer are dispersed on the x-axis, and layers are stacked onto each other following the z-axis. The lattice representation is summarized into a Form string as shown on the top. The Form describes each SSE by the layer, relative position in the layer, and secondary structure type separated by a dot (N- to C-terminal sequence order is preserved). **B:** A multi-Form string created by assigning some SSEs as mandatory and others as optional. The flexibility allows the sampling of a range of architectures and topologies. **C:** Comparing the exploratory capacity between a simple Form (5 SSEs), a multi-Form (a minimum of 5 and a maximum of 10 SSEs) and the known space of protein folds (as classified by CATH). The simple Form nearly samples as many existing topologies as known, while the multi-Form greatly generates more topologies than what can be found in nature.

## Results

Evolution proceeds incrementally through a random and sparse sampling of the possible sequence space which in turn populates the protein structure space. However, nature seems to show a tendency to reuse the same protein structures repeatedly based on the observation that the discovery of new protein folds has become rare [34]–[36]. In some regards, the mapping of structural space poses a number of challenges since it depends on the structural definition and coarseness of the structures. For a more systematical exploration of the structural space, Taylor and colleagues defined an idealized SSE lattice representation that can easily be captured through a simple string descriptor called Forms [36], [37] (Fig. 1A). The Form parametrization describes proteins as layered topologies, with each layer being composed of a defined number of either α-helices or hydrogen bonded β-strands (Fig. 1A). Although constrained by its grid-like tabular system, a wide range of structural configurations can systematically be defined, potentially allowing to fully explore a protein topological space at orders of magnitude larger than the natural space currently characterized (Fig. 1B, 1C). Although Forms are well suited for protein topology comparison and classification [39], using them for *de novo* design is challenging due to the loss of crucial structural and sequence features, including native tertiary configurations of SSEs and sidechain representations.

### Sketching native-like protein backbones from Forms

Given the Form description, the TopoBuilder starts by placing ideal SSEs at their respective relative positions as specified by the Form description, creating a three-dimensional (3D) backbone object containing only SSEs, which we refer to as “Sketch”. We define the layer stacking along the z-axis (Fig. 2A), with inter-layer separations of 8 Å for β-sheets, 10-11 Å for α-helices, and mixed structures [40]. The y-axis aligns with the directionality of the SSEs (Fig. 2A), and the intralayer spacing between adjacent SSEs is defined along the x-axis and typically of 10 Å for α-helices and 4.85 Å for β-strands [41].

**Figure 2.**
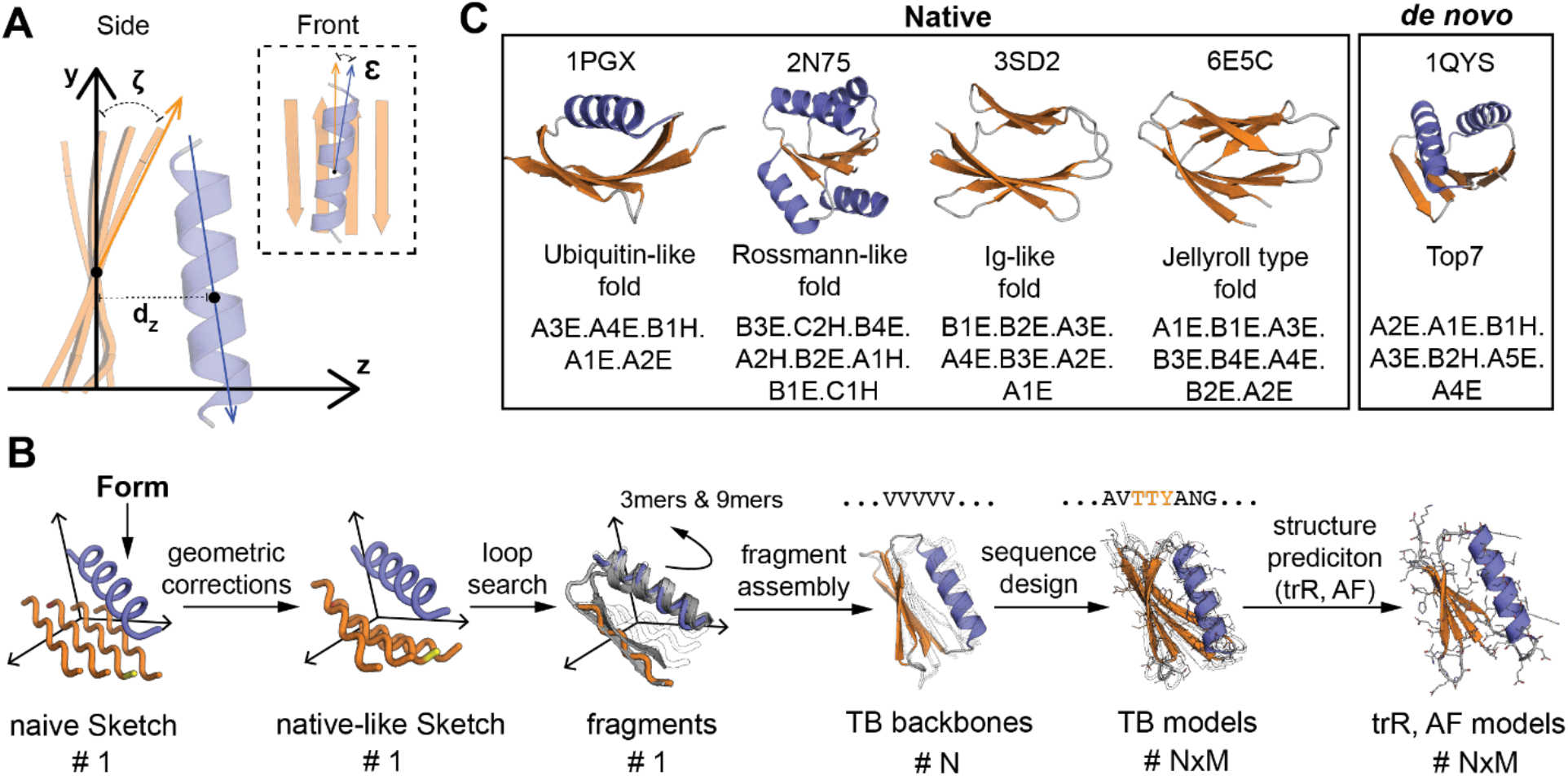
Setup for geometric parameterization and correction in example protein folds. **A:** Example of four geometrical correction parameters calculated from the matches found by MASTER and exemplified on a Sketch. **B:** The generic TopoBuilder pipeline starting from a Form that expands into a naive Sketch, which is then corrected into a native-like Sketch. Loops are searched to connect the SSEs and fragments of size 3 and 9 are created (3mers and 9mers). Using the fragments and distance constraints from the native-like Sketch, N polyvaline backbones are assembled and NxM sequences designed. **C:** Four examples of native protein structures and a *de novo* protein with their corresponding Form description covering a variety of structural folds.

The Sketch representation does not contain SSE connecting loops and has no sequence information. Furthermore, the naive Sketch presents a rather non-native configuration of the SSEs composing the protein structure strongly hinting towards the possibility that such structures will likely present non-designable configurations. Fine-grained structural details at the secondary structure and tertiary levels such as β-sheet pleating and curvature, and α-helical packing are absent. These features are difficult to sample automatically and correctly with low-resolution scoring functions.

We hypothesized that *grosso modo* parametric corrections per SSE could incorporate global native structural features and improve the Sketches’ designability (Fig. 2A and Supp. Fig S2A-E). To do so, we implemented a module that computes on-the-fly geometrical statistical corrections for each SSE from native structures (see Methods). Briefly, the Sketch is first divided into two-layer components based on adjacent layers. These sub-structures are then iteratively queried against a database of natural protein structures using the software MASTER [42], [43] and structural geometry parameters calculated from the retrieved matches and used to correct the relative positioning of the SSEs in the Sketch. The iterative nature of the process along the layers results in a hierarchical refining procedure where the previously corrected sub-structures help to contextualize the correction in the next layer for coherent native-like SSE placements (Supp. Fig. S3).

A small set of parametric corrections includes two rotational parameters and one translational parameter per SSE (Fig. 2A and Supp. Fig. S2A-E). We compute the twist angle (ζ) between the vector pointing along the length of the SSE and the plane spanned by the layer. Between two adjacent layers, we express the angle (ε) as the shear between the layer planes, and the inter-layer distance (d_z_) as the distance from one layer to the next one. The individual parameters shift the SSEs from a naive configuration towards a native arrangement. For example, a β-sheet in the naive configuration is “flat” because of the initial placement of the SSEs that are fully ideal and aligned next to each other. The natural occurring twist within β-sheets can be approximated by twist angles ζ [38]. Hence, the geometric corrections attempt to optimize the global arrangement of the SSEs and generate topologies with native-like features.

### Backbone assembly and sequence design of native-like Sketches

To obtain sequence-designed structures, the native-like Sketches are subjected to several steps: (1) loop building to connect the SSEs; (2) structural diversification starting from the initial native-like Sketch; (3) sequence sampling and selection of best scoring designs. All these steps were performed using tools provided by the Rosetta software suite [44], and more details are given below and in the Methods section (Fig. 2B and Supp. Fig. S1A-F, 3, 4).

To build fully connected structural templates, we query native loop segments that can bridge the gaps between the SSEs that compose the Sketch (see Methods). We avoid time-consuming computational loop closure algorithms by leveraging structural information of the retrieved loops and generating structural fragments (3-mers and 9-mers) (ABEGO torsions) (Supp. Fig. S4). We use the previously developed Rosetta FunFolDes (FFD) [45], [46] protocol to generate an initial set of poly-valine backbone conformations using Cα-based distance restraints calculated between all different SSEs and perform sequence design. The structural fragments impose native backbone signatures at the local level, while the overall topology is tightly controlled by the distance restraints. We modified the Rosetta energy function at every stage of the folding simulation to include hydrogen bonding- and SSE pairing terms favoring the correct pairing between β-strands [47]. Each assembled backbone is fitted with a set of optimal sequences via the Rosetta FastDesign protocol [44]. During this stage, AA sampling restrictions per position were added, such as layer definitions (core, surface, or boundary, profiles from structural fragments) [48], [49], and secondary structure type assignments (α-helix, loop, or β-strand). A bonus term enhancing secondary structure formation at defined positions was included in the energy function at the desired protein segments [50].

### *De novo* design of five protein folds

To showcase the TopoBuilder *de novo* design framework and assess its performance, we attempted to *de novo* design five folds of distinct structural complexities (Fig. 2C). We selected four native folds: a 2-layered α/β Ubiquitin-like fold, a 2-layered β-sandwich Ig-like fold, a 2-layered β-sandwich Jelly-roll, and a 3-layered α/β/α Rossmann-like fold. In order to investigate the generalization capability of the framework to the space of novel folds, we included a 2-layered α/β Top7-like fold. Of note, the Top7 structure (PDB 1QYS) was excluded from all databases used during the correction searches to avoid any biases coming from the solved structure. The Ubiquitin-like fold is composed of a α-helix packed onto a 4-stranded β-sheet. Both terminal β-strands pair in a parallel direction and are located in the center of the sheet making non-local hydrogen bond contacts, while the edge strands form a β-α-β motif. The Jelly-roll and the Ig-like have both non-local β-β motifs. Our drafted Jelly-roll has three β-arcade motifs and the Ig-like fold is made from a β-arcade on one and a long β-arch on the other side. The architecture of the intended Rossmann fold contains a 4-stranded central β-sheet that is flanked by two helices on the top and two helices at the bottom and can be decomposed into three interlocked β-α-β motifs. Lastly, the Top7 fold is defined by two interlocked β-α-β motifs with two additional terminal core strands. Each of the folds has its unique complexity with specific tertiary motifs that need to be arranged realistically with the correct geometries. Especially non-local interactions and connections have been difficult to build and design, and multiple detailed analyses were needed to define effective design rules for single folds and domains [26]–[28], [51], [52].

For the generation of the backbones, we used default SSE lengths of 5 to 8 residues for β-strands, 17 residues for long α-helices, and 13 residues for short α-helices. Thus, we did not employ SSE lengths of specific native examples but rather used these as a rough guide for the overall topology. To probe the impact and contribution of the parametric corrections, we first performed baseline design simulations guided only by an uncorrected Sketch that we refer to as “naive Sketch”. We then corrected the Sketch using several combinations of parameters (termed “native-like Sketch”) to assess their importance and find a minimal and optimal parameter combination. In the first scenario, we solely used the twist angle ζ correction, the second scenario consisted of the corrections ζ + d_z_, and the third scenario simulated with ζ + d_z_ + ε (Supp. Fig. S5, S6, S7). For each of the different scenarios, a total of 1,000 decoys were generated.

### Corrections induce native features in idealized folds

For the native folds, the corrections were derived from a set of ~15-25 structurally distinct proteins (Supp. Fig. S8A-C). The Top7 fold-derived Sketch had a lower number of matches that were extracted from larger protein domains with similar SSE dispositions and connectivities (Supp. Fig. S8A-C). We compared the native-like and the naive decoys for each of the five selected folds (Fig. 3A and Supp. Fig. S9A-E). The ζ angle distributions (Fig. 3B) show that native-like decoys have a twist in β-strands and native side-to-side configurations for α-helices, while the distributions of the naive decoys retain a ζ angle around 0° indicating that no twist was induced in the strands during the fragment assembly folding simulations. Similarly, the ε angle (Fig. 3C) and the d_z_ distance (Fig. 3D) that improve the layer packing geometry, follow the native distributions for the native-like but not for the naive Sketch-derived decoys, showing that fragment insertion protocols are insufficient to correct global topological features. Also, a relaxation without restraints after the folding and design simulations did not yield native geometries in naive decoys, suggesting that geometric corrections were necessary to guide the folding trajectories of the native-like sketches.

**Figure 3.**
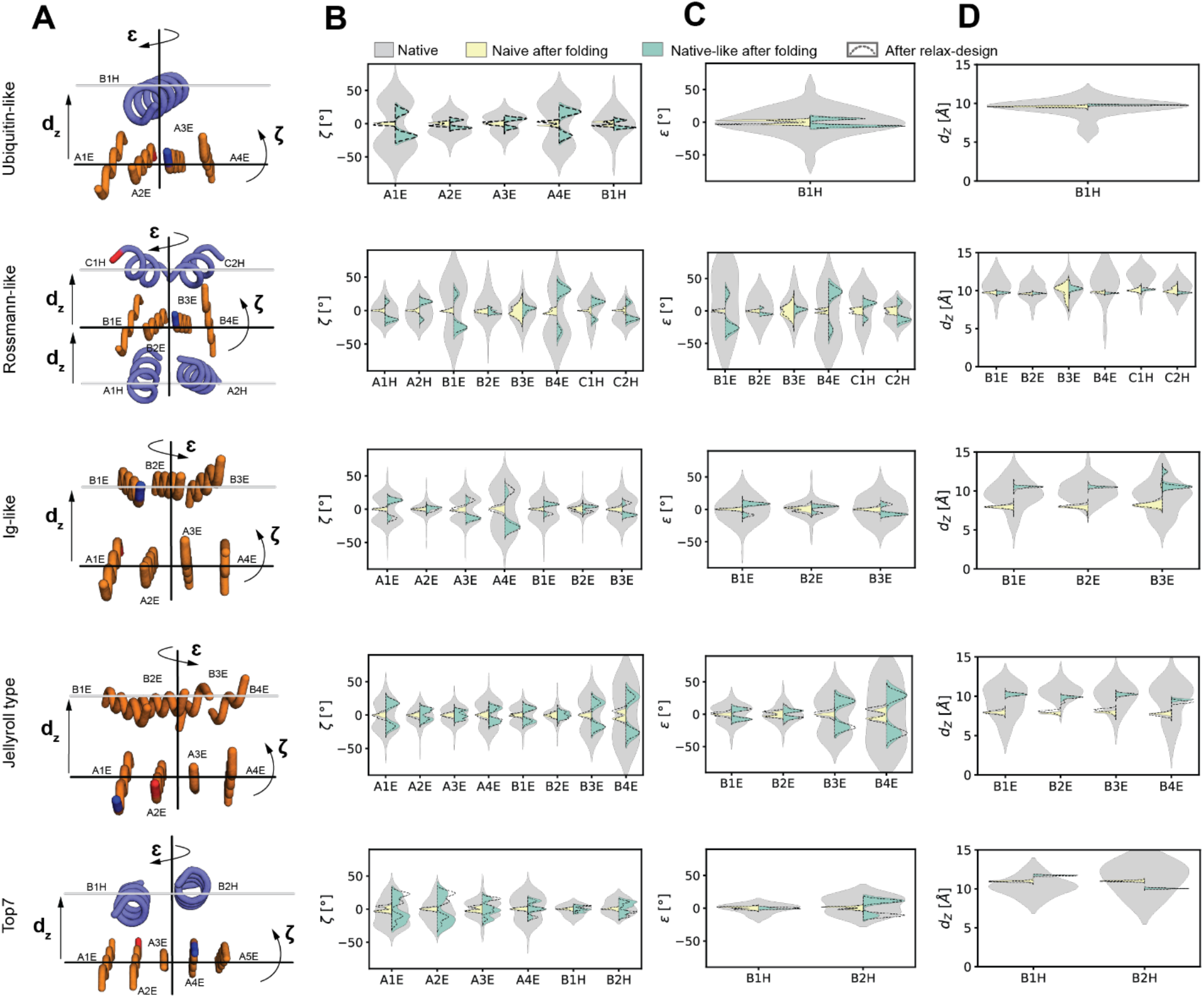
Geometric corrections improve topological features of the designed structures. **A:** The native-like Sketches of the example folds with three of the geometric parameters indicated. **B:** Distributions of the ζ correction parameter computed from the retrieved matches. The ζ correction parameter captures features related to the orientation of the different layers. **C:** Distributions of ε correction parameter computed from the retrieved matches. The ε correction parameter captures features related to the orientation of the different layers. **D:** Distributions of the d_z_ correction parameter computed from the retrieved matches. The d_z_ correction parameter captures the distance between layers in the Sketch. **B,C,D:** The native geometry distributions are shown in gray. In yellow, the output of simulations (1,000 decoys) using a naive Sketch without correction of the SSEs, which tends to result in flat β-sheets and α-helical stackings. Shown in green are simulations (1,000 decoys) derived from native-like Sketches where the outputs more closely follow the native distributions.

### Assessing designability through sequence-to-structure compatibility

To assess the difference between sequences from the naive and native-like designs, we first used BLASTp [53] to search for similar sequences in the natural repertoire. However, the few hits (Evalue < 0.01) found did not match the target fold showing that there were no evident fold signatures in either of the design sets (Supp. Fig. S10A,B). Therefore, we used two orthogonal deep learning protein structure prediction engines trRosetta (trR) [54] and AlphaFold (AF) [55] to predict structural models for all designed sequences (without MSA generation, i.e. in single-sequence input mode). We computed the template modeling (TM)-scores and root-mean-square deviations (RMSDs) between the TopoBuilder designs and the predicted structures by trR and AF (Fig 4A, 4B, Supp. Fig. S11A,B). We hypothesized that, if our native-like backbones have improved designability, they could lead to sequences with stronger propensities for the respective fold and consequently to more accurate structure predictions in contrast to the naive Sketch-derived designs.

**Figure 4.**
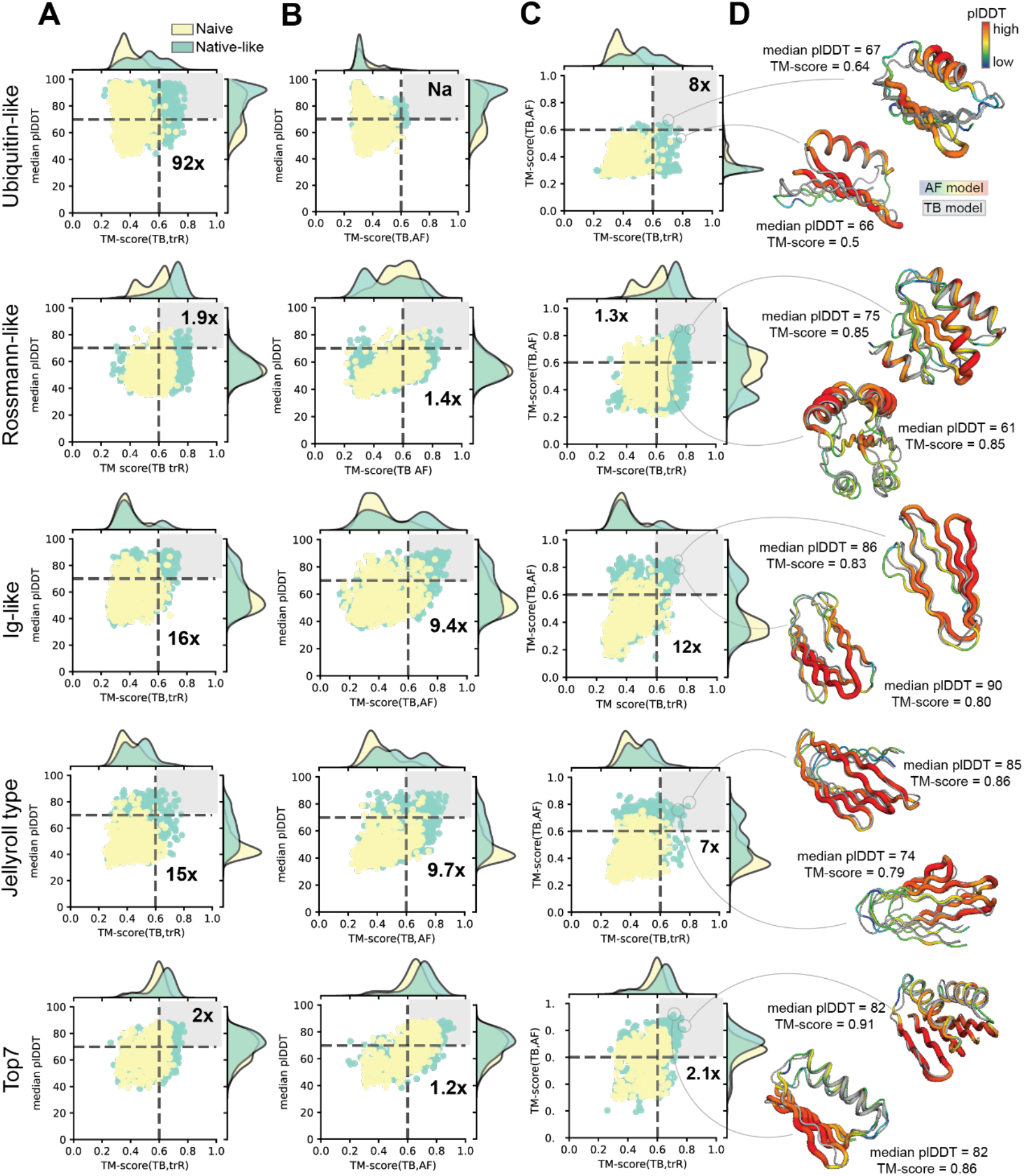
*In silico* assessment of the sequence quality of the TopoBuilder designs. Evaluation of sequence generation improvements when comparing the usage of naive and native-like Sketches. Boost in quality for native-like derived sequences with m sequences in the upper right quadrant (best sequences according to TM-score to the target structures and median confidence scores from AF (plDDT)). **A,B,C:** The plDDT threshold is fixed to 70, and the TM-score is fixed to 0.6 in order to select the respective double-positive populations. **A:** The plDDT scores versus the TM-scores computed from the TB models aligned onto trRosetta models. **B:** The plDDT scores versus the TM-scores computed from the TB models aligned onto AF models. “Na” labels an infinite enrichment ratio given that in the naive Sketch guided designs there were no sequences that fulfilled the defined thresholds. **C:** Comparison of the TM-scores computed between the TB models and AF or trR models. **D:** Examples of TB native-like designs. In color the AF model according to its confidence and in gray the TB model.

The comparisons for the naive and native-like design sets in terms of accuracies for the structure prediction simulations, do not show clear differences. For the naive designs, we observed low RMSDs at around ~2 Å trR and ~1.8 Å AF while the native designs achieved RMSDs of ~1.5 Å and ~1.2 Å for trR and AF, respectively (Supp. Fig. S12A). Similarly, for TM-scores the naive design sets peak at ~0.7 for trR and ~0.8 for AF, and the native-like reached TM-scores of ~0.8 for trR and ~0.9 for AF (Supp. Fig. S12B).

Despite the lack of clear preference at the level of individual design metrics, we noticed that comparing the naive- and the native-like derived sequences based on two metrics shows population differences (Fig. 4). We compare the AF predicted local Distance Difference Test (plDDT) versus the TM-score between the TopoBuilder (TB) and the trR models (TM-score(TB,trR)) or AF models (TM-score(TB,AF)).

To quantify the upper right quadrant (double-positive) population, we set the plDDT threshold to a minimum of 70 and adjust the TM-score thresholds to 0.6. Analyzing the population difference of the double positives e.g., sequences with plDDT > 70 and TM-score(TB,trR) > 0.6 shows an enrichment of the native-like designs ranging from 1.9x in the Rossman-fold designs to 92x in the Ubiquitin-like designs (Fig. 4A). Similarly, we then compared the plDDT against the TM-score between the TB and the AF models (TM-score(TB,AF)) and observed similar enrichments ranging from 1.2x for the Top7 designs to 9.7x for the Jellyroll designs (Fig. 4B). Lastly, we compared the TM-score(TB,AF) versus the TM-score(TB,trR) to assess the agreement between the two prediction methods fixing the TM-score(TB,AF) and TM-score(TB,trR) thresholds to 0.6. We see a clear enhancement of the native-derived double positives ranging from 1.3x for the Rossmann designs to 12x for the Ig-like designs (Fig. 4C).

To further assess the increased performance induced by the corrections, we projected the three score-pairs (1) plDDT and TM-score(TB,trR), (2) plDDT and TM-score(TB,AF), and (3) TM-score(TB,trR) and TM-score(TB,AF)) onto their respective diagonal and computed the receiver operating characteristic (ROC)-curve and the area under the curve (AUC) (Supp. Fig. S13A,B,C). The ROC-AUC indicates the degree of separation between the naive- and native-derived projected distributions independent of an arbitrary set threshold. Most ROC-AUC values are in the range of 0.63 - 0.7 across the three different score pairs additionally showing that our corrections improved the *de novo* design of proteins (Supp. Fig. S13A,B,C).

The superior structural metrics produced by trR and AF for the sequences generated on native-like TB backbones suggest that these sequence improvements arise from more designable backbones with native-like features.

### Experimental validation of novel sequences

We next sought to experimentally test whether the TB designs were folded and stable in solution. We investigated the top TB models by TM-score(TB,trR), obtained the synthetic genes for 54 designs, and expressed and purified them from *E. coli* (see Methods). From the tested designs, 3 Ubiquitin-like, 4 Rossmann-like, 3 Ig-like, 1 Jelly-roll type, and 2 Top7 like fold proteins expressed soluble. A total of 2 Rossmann designs, 1 Ig-like fold designs, and 2 Top7 designs (Fig. 5A,B) had size exclusion chromatography (SEC-MALS) peaks (Fig. 5C) with an apparent molecular weights of monomers or small oligomeric species. Corresponding monomeric or small oligomeric species (dimer or trimer) were examined by circular dichroism (CD) spectroscopy (Fig 5D). In all cases, the CD spectra were consistent with the respective target structures, with the characteristic profiles of α/β and mainly β proteins. The designs were thermostable with melting temperatures above 90 °C (Supp. Fig. S14).

**Figure 5.**
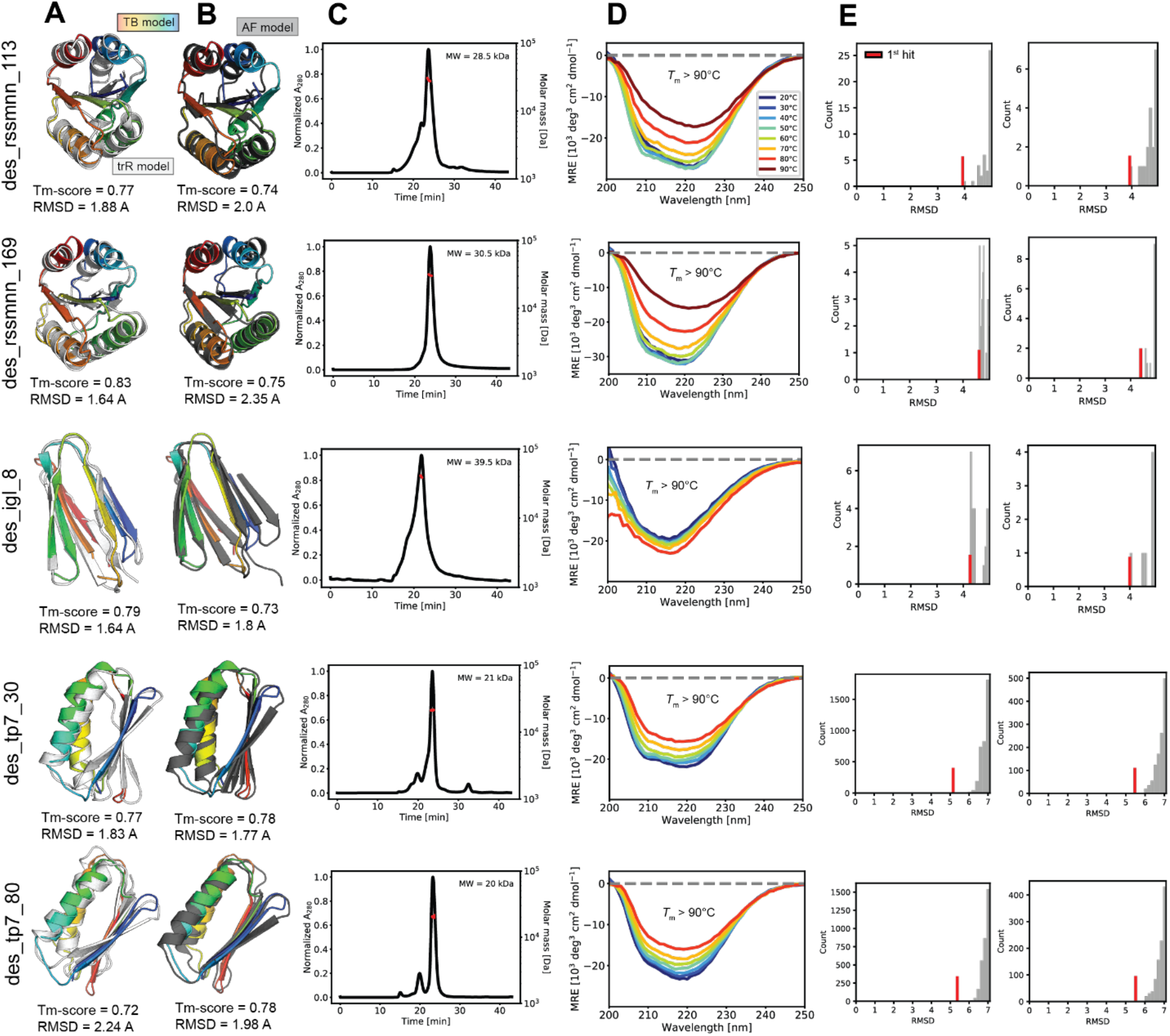
Experimental characterization of native-like TopoBuilder designs. **A:** TopoBuilder (TB) and trRosetta (trR) models superimposed. **B:** TB and AlphaFold (AF) models superimposed. **C:** SEC-MALS elution profiles showing the oligomeric state of the designs in solution. The TB designs des_rssmnn_113, des_rssmnn_169, des_tp7_30 and des_tp7_80 show a dimeric while the des_igl_8 shows a quaternary state in solution. Red dots show the molecular weight as determined by SEC-MALS. **D:** Circular dichroism spectroscopy with thermal denaturation. The TB designs adopt folded structures in solution. The melting temperatures (*T*_m_) were obtained by fitting to the denaturation curves shown in Supp. Fig. S14. E: Fast RMSD-based structure searches (PDB and AF databases) showed that the closest structures found in existing databases were typically below 5 Å.

As a retrospective analysis of the experimentally characterized sequences, we performed a deeper analysis to the predictions performed by trR (Fig. 5A) and AF (Fig. 5B). The five designs (Rossman, Ig-like, Top7) recapitulated the target structures accurately, strongly indicating that our designs folded in the desired conformation. To evaluate the structural similarity relative to the natural repertoire, we queried the PDB [56] and the AF [57] (Fig. 5E) databases for similar protein structures. While for both natural folds (Rossmann-like and Ig-like) we found first hits at ~4 Å RMSD, the Top7 designs are far away from any natural protein folds with the first hit at ~5.5 Å RMSD.

To reveal potential underlying sequence clusters, we used the found structural matches from the PDB and AF database and performed pairwise alignments. We then calculated the Blosum62 distances for each alignment (i.e. the sum of each individual Blosum62 score), and clustered them hierarchically (Supp. Fig. S15A and Supp. Fig. S16A). Similarly, to categorize the conformations we calculated pairwise RMSDs followed by a hierarchical clustering (Supp. Fig. S15B and Supp. Fig. S16B). Our designs are generally well integrated within the hierarchical cluster trees showing native compatibility. To search for close members in sequence and structure space jointly, we gathered the sequence and structure features and projected the data into two dimensions through a principal component analysis (PCA) (Supp. Fig. S15C and Supp. Fig. S16C). We observe that our designs are close to native clusters, further indicating their sequence and structure nativeness. For the native folds, the matches are of the same fold family. Interestingly, des_rssmnn_113 has a *de novo* designed Rossmann fold (PDB 2LV8) and natural Rossmann domains (PDB 1MZP and 4IZ6) as cluster members, hence the design likely incorporated general native sequence- and structure features. The des_tr7_30 based on the Top7 *de novo* designed fold has native cluster members that structurally fit well, but have different connectivities (e.g., PDB 6NR1, 4QTP or 4QDJ).

Taken together, the data indicate that five designs across three different folds designed using native-like templates adopted stable monomeric or dimeric states with the expected secondary structure content and accurate structural predictions by AF.

## Discussion

The TopoBuilder *de novo* design method enables the generation of artificial proteins from a minimal string description (Form). The Form description drafts the overall target topology and enables a fast and systematic fold-space exploration [39], [58]. Combined with the TopoBuilder *de novo* design framework, virtually any protein Form description can be constructed and designed.

Our computational and experimental assessments show that geometrical corrections inferred from native structural sub-motifs that compose the folds provide enough information to improve the designability of protein backbones [33]. When analyzing the designed sequences with state-of-the-art structure prediction tools, we identified a larger fraction of successfully recovered structures from sequences derived from corrected (native-like) backbones than those from naive backbones. The experimental characterization of multiple designs shows that the TopoBuilder *de novo* design framework generates realistic designs that adopt the target fold and are thermodynamically stable.

Ultimately, our analysis shows that the current scoring functions and fragment assembly methods are insufficient for the generation of designable backbones without the guidance of natively arranged SSEs. Many of the *de novo* protein design rules rely on the generation of structured loops to guide the SSEs’ placements. Here, we present an alternative and complementary solution that is fully automated upon the definition of the length of the SSE. Instead of focusing on structured loops, we optimize and correct the global placements of SSEs and thereby implicitly guide the loop geometries.

Our strategy should further enable *de novo* design to non-experts, improve and streamline future protein design efforts. The insights we gained from the parameters for a variety of complex fold examples can be harnessed and support the future discovery and understanding of protein architectural principles. Our work also opens new possibilities for computational protein designers that may want to design for function such as the scaffolding of functional proteins via incorporating known or predicted functional sites, or protein assemblies with *de novo* designed domains.

## Material and Methods

### Computation of number of architectures and topologies from Forms

To approximate the number of architectures and topologies from a Form description (Fig. 1C), we treated all SSEs elements as of the same type. Thus, if a topology consists of 5 SSE elements (n = 5), we did not differ between architectural and topological variations on the number of helical (H) and strand (E). We related the layered architectures to free polyominos without holes (to discard underpacked architectures from the count) which can be found under A000104 (https://oeis.org/A000104). This enabled a simple and fast lookup of the number of architectures for a Form with n SSEs. The total number of topologies for one architecture can be computed through n!.

### Topological refiner

Structural subunits within a particular protein Sketch are queried using MASTER against a database of native proteins. The database consisted of structures of 70-250 residues. To speed up the matching procedure, we only searched over fold-relevant structures by filtering the database based SSE content and structural features, e.g. for Ig-like and jelly-roll folds we only quired β-sandwich architectures, and for Ubiquitin-like, Rossmann-like and Top7 folds we quired 2 or 3 layer α/β architectures. The RMSD thresholds for the selection of matches ranged from 2-3 Å. We processed each MASTER match by fitting a vector along each SSE (Supp. Fig. S2A,B,C) via performing a Principal Component Analysis (PCA, for more details see [37]) over all Cα atoms within the SSE. Naturally, the first (major) eigenvector returned by the PCA points along the SSE length.

For each full layer (including all SSEs) we compute the first three eigenvectors (Supp. Fig. S2D). This will result in a local coordinate system for the layer where the first eigenvector is along the y-axis (along the lengths of the SSE), the second eigenvector the x-axis (towards the side), and the third eigenvector the z-axis. The first eigenplane defines the 1-2 (major-side) plane that is formed by the first two eigenvectors. The 1-3 (major-perpendicular) plane generated by the first and third eigenvector splices the layer along the length of the SSE in half. Lastly, the 2-3 (perpendicular-side) plane is computed using the second and third eigenvector and halfs the layer roughly along the center of each SSE. Having abstracted from atoms to simple geometric objects such as vectors and planes, multiple parameters can be efficiently computed (Fig. 2A). Considering two adjacent layers, one can compute several geometric features. Here, we use the ζ angles, which are the angles between the first SSE eigenvectors and the layer (1-2) plane. The ε angles, which are the shear angles between the two layers, can be calculated as the mean across all first SSE eigenvectors with the corresponding 1-3 eigenplane. The interlayer distance d_z_ can be computed as the mean across all SSE center distances to adjacent layers. Lastly, the sheer distances are calculated as the distances between the 2-3 planes to the SSE centers.

### Loop Assembler

To connect the SSEs within the Sketches we developed a workflow to harness structural features from secondary structure connecting loops in native proteins (Supp. Fig. S4). For each gap, we performed independent MASTER searches with their corresponding two SSEs. The algorithm iteratively matched two consecutive SSEs against a database of protein structures. For each gap, matches with an RMSD smaller than 3.2 Å were kept and clustered based on their loop lengths (the maximum length allowed was 7 residues), and the most populated cluster lengths were selected. Subsequently, the selected loops were filtered with respect to their ABEGO torsion profiles [59]. Loops displaying the same ABEGO dihedral angle for each residue were removed from the set, leaving a single loop per ABEGO profile. Structural fragments of sizes 3 and 9 (3mers and 9mers) were generated for the loops and SSE alignment regions in agreement with the ABEGO distributions observed in the loop matches. These fragments were then used in subsequent fragment assembly steps to generate fully connected folded structures. With this strategy we avoided performing explicit loop building and closure sampling to add the loops on the Sketch as this requires computationally expensive structural sampling without the guarantee of closing the gap effectively.

### Folding of poly-valine backbones from Sketches

Structural backbones were generated using the Rosetta FunFolDes (FFD, NubInitioMover) [47] fragment assembly folding simulations. The 3mers and 9mers structural fragments derived from the loop assembler were used for the folding simulations introducing local native-like interactions and structural patterns in the loop regions. To control the global conformation, Cα-Cα distance restraints between all SSEs of the Sketch were used during the folding trajectories. We modified the energy function during the four stages of the folding simulations by including short- and long-range hydrogen bonding terms and SSE formation enhancing terms to yield compact backbones with favorable non-local interactions and global tertiary geometries. We also imposed secondary structure assignments to each residue to bias the simulations that were derived from the secondary structure definition of the Sketch.

### Sequence design and structural relaxation of backbones

We used the Rosetta software to design sequences for each of the generated poly-valine backbones followed by structural relaxations [44]. The sequence design operations at each residue position were restricted through (1) SSE propensity calculated from the Sketch, (2) “layer” assignment calculated from the backbone conformations where the “core”-, “boundary”-, and “surface” layers are defined through the number of Cβ neighbors, and (3) an AA sequence profile derived from the structural fragments to upweight frequent AAs. The structural relaxations were performed under long-range Cα-Cα distance restraints derived from the backbone conformations. Additionally, we added a topological bonus term to the energy function to favor conformational changes with corrected strand- and helix pairings and helix-sheet packing. The topological restraints were extracted from the Sketch. A total of 200 poly-valine backbones were generated and 5 sequences designed for each backbone. This led to a total of 1,000 different sequences per Sketch.

### Structure predictions using trRosetta and AlphaFold

We used the trRosetta (trR) [54] and AlphaFold (AF) [55] deep neural networks to predict structural models from the designed sequences. We predicted structural models for all sequences (1,000 per Sketch) using trR and AF in single sequence mode e.g., omitting the time-consuming step of generating multiple-sequence alignments (MSA). Additionally, we only generated a single AF model (instead of five), making the computation 5x faster. All inference calculations were parallelized onto 1,000 CPU cores.

We aligned each trR and AF model onto the respective TopoBuilder (TB) model using the template modeling (TM) alignment algorithm [60]. The algorithm iteratively optimizes the superposition of segments with similar local structures. The superposition between the two structural models is evaluated through the TM-score, a measure of the distance between Cα atoms of aligned residues in target and template, normalized by protein length. TM-align returns the TM-score and a best-fit RMSD.

### Protein Expression and Purification

The 54 best designs by TM-score(TB,trR) were selected for experimental validation. DNA sequences of the designs were purchased from Twist Bioscience. For bacterial expression, the DNA fragments were cloned via Gibson cloning into a pET11b followed by a terminal His-tag and transformed into Escherichia coli BL21(DE3). Expression was conducted in Terrific Broth supplemented with ampicillin (100 μg/ml). Cultures were inoculated at an optical density (OD) 600 of 0.1 from an overnight culture and incubated in a shaker at 37 °C and 220 r.p.m.. After reaching an OD600 of 0.6, expression was induced by the addition of 0.4 mM IPTG and cells were further incubated overnight at 20 °C. Cells were harvested by centrifugation and pellets were resuspended in lysis buffer (50 mM TRIS, pH 7.5, 500 mM NaCl, 5% glycerol, 1 mg/ml lysozyme, 1 mM PMSF, 4 μg/ml DNase). Resuspended cells were sonicated and clarified by centrifugation. Ni-NTA purification of sterile-filtered (0.22 μm) supernatant was performed using a 5 ml His-Trap FF column on an ÄKTA pure system (GE Healthcare). Bound proteins were eluted using an imidazole concentration of 500 mM. Concentrated proteins were further purified by size exclusion chromatography on a Hiload 16/600 Superdex 75 pg column (GE Healthcare) using PBS buffer (pH 7.4) as mobile phase.

### Circular dichroism spectroscopy

Far-UV circular dichroism spectra were collected between wavelengths of 190 and 250 nm on a Jasco J-815 circular dichroism spectrometer in a 1mm path-length quartz cuvette. Proteins were diluted in 10 mM Phosphate-buffered saline (PBS) at concentrations between 20 and 40 μM. Wavelength spectra were averaged from two scans with a scanning speed of 20 nm/min and a response time of 0.125 s. The thermal denaturation curves were collected by measuring the change in ellipticity at 220 nm from 20 to 90 °C with 2 or 5 °C increments.

### Size-exclusion chromatography combined with multi-angle light scattering

Multi-angle light scattering was used to assess the monodispersity and molecular weight of the proteins. Samples containing 80–100 μg of protein in PBS buffer (pH 7.4) were injected into a Superdex 75 10/300 GL column (GE Healthcare) using an HPLC system (Ultimate 3000, Thermo Scientific) at a flow rate of 0.5 ml/min coupled in-line to a multi-angle light-scattering device (miniDAWN TREOS, Wyatt). Static light-scattering signal was recorded from three different scattering angles. The scatter data were analyzed by ASTRA software (version 6.1, Wyatt).

## Acknowledgments

We thank the members of the Protein Design and Immunoengineering group (LPDI, EPFL, Lausanne) for helpful discussions. We thank Arne Schneuing for the critical reading of the manuscript. We would like to thank EPFLs Scientific IT and Application Support Center for their support on the computational infrastructure. We would like to thank the Protein Production and Structure Core facility at EPFL for their support on the protein biophysical characterization experiments. B.E.C. is a grantee from the European Research Council (Starting grant - 716058), the Swiss National Science Foundation, and the Biltema Foundation. Parts of the computational simulations were performed at the CSCS - Swiss National Supercomputing Centre through a grant obtained by B.E.C.. Z.H., and S.R. are supported by a grant from the National Center of Competence in Research in Chemical Biology.

## Additional information

### Author information

These authors contributed equally: Zander Harteveld, Jaume Bonet.

### Contributions

B.E.C. conceived the initial idea and refined it together with Z.H., J.B., F.S., C.Y., and B.E.C.; Z.H., S.R., J.B., and B.E.C designed and performed experiments. Z.H. and J.B. wrote the software. S.R. purified the designs and performed protein-biochemical characterization. Z.H. performed in silico structural analysis and modeling with the support of F.S. and C.Y.; B.E.C. directed the work. B.E.C. and Z.H. wrote the manuscript with support from all authors.

## Supplementary figures

**Supplementary Figure S1.**
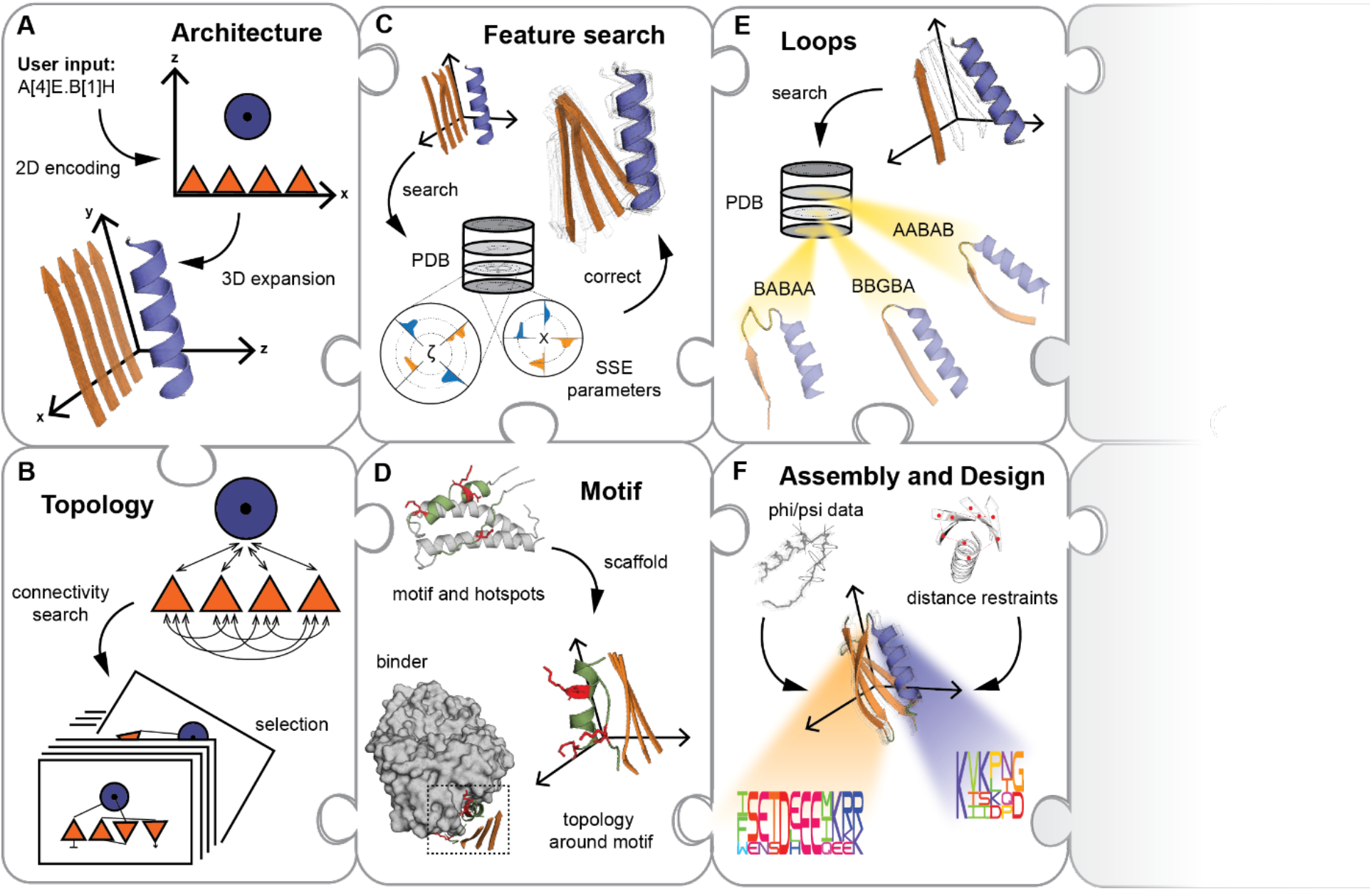
Overview of the modules in the TopoBuilder *de novo* design framework. The TopoBuilder *de novo* design framework can easily be extended and modified by adding custom modules with the provided Python base-code. **A:** The generic pipeline takes a Form string as input and generates a 3D expansion of the architecture or topology Sketch. **B:** If the topology is not specified, all possible topologies are generated for the user to select a single topology. **C:** At the heart of the pipeline, the topological refiner protocol searches a database of natural protein structures for geometric features to recover a native-like configuration of the Sketch. **D:** An additional module enables the incorporation of structural motifs for the design of function. **E:** The loops are reconstructed implicitly via first searching for natural loops that are suited to bridge the SSE gaps and secondly creating structural fragments from the identified loop regions. **F:** The structural model built via Rosetta fragment assembly (Rosetta FunFolDes) and designed using the FastDesign method.

**Supplementary Figure S2.**
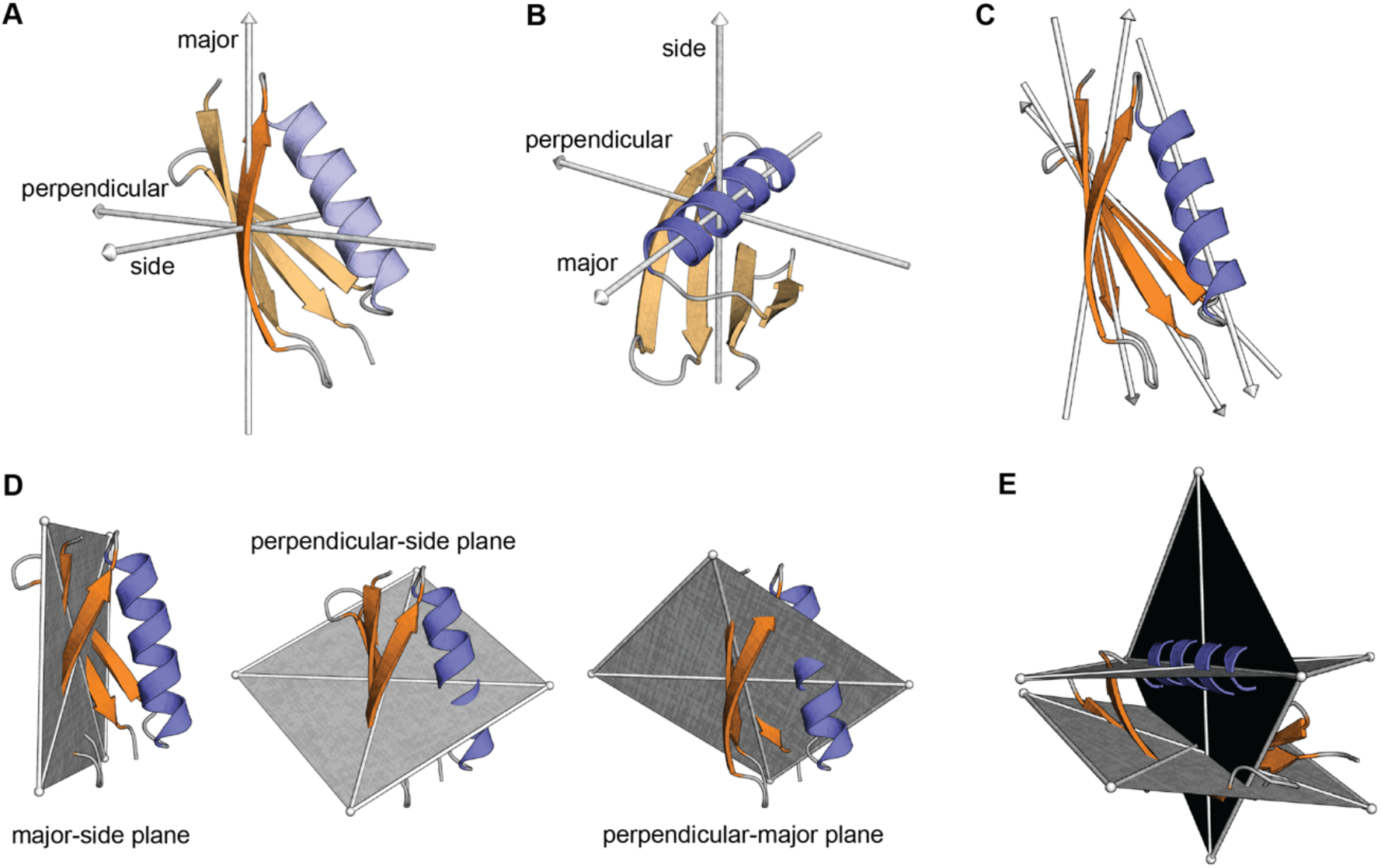
Example of geometric parametrization performed by the TopoBuilder. **A**: Eigenvectors computed for a single strand of a Ubiquitin-like fold structure (PDBID 1PGX). The major eigenvector points along the major axis of the SSE. **B**: The eigenvectors computed for a single helix of the 1PGX structure. The major eigenvector aligns with the axis along with the helical structure. **C**: All major eigenvectors for each of the SSE of the 1PGX structure. The positions and directions of the SSE can be described by the major eigenvectors. **D**: The set of the eigenplanes computed for the sheet (layer A) 1PGX structure. **E**: The major-side planes for the sheet and the helix of the 1PGX structure.

**Supplementary Figure S3.**
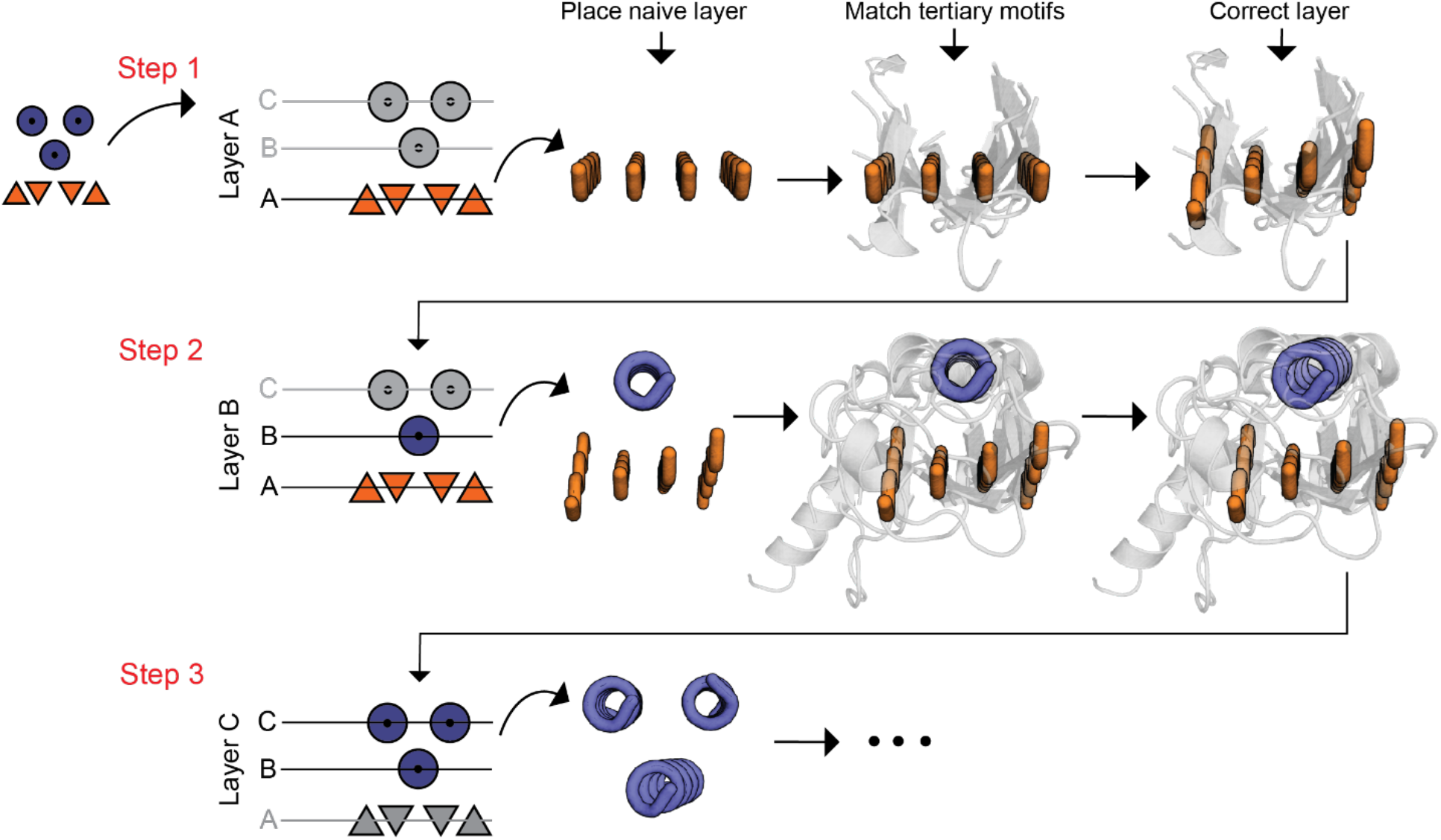
Topological corrections performed by the TopoBuilder through a hierarchical refinement procedure. The topological refiner module starts at layer A (single layer) and searches for similar tertiary motifs to compute geometric corrections. The corrections are then used to generate a native-like version of layer A. The second step integrates the native-like layer A and places the next layer (in this example a helix) on top, performing the structural search and correction calculations that are then applied to layer B. The third step places layer C naively onto the corrected layer B and repeats the correction procedure in the presence of the corrected layers A and B.

**Supplementary Figure S4.**
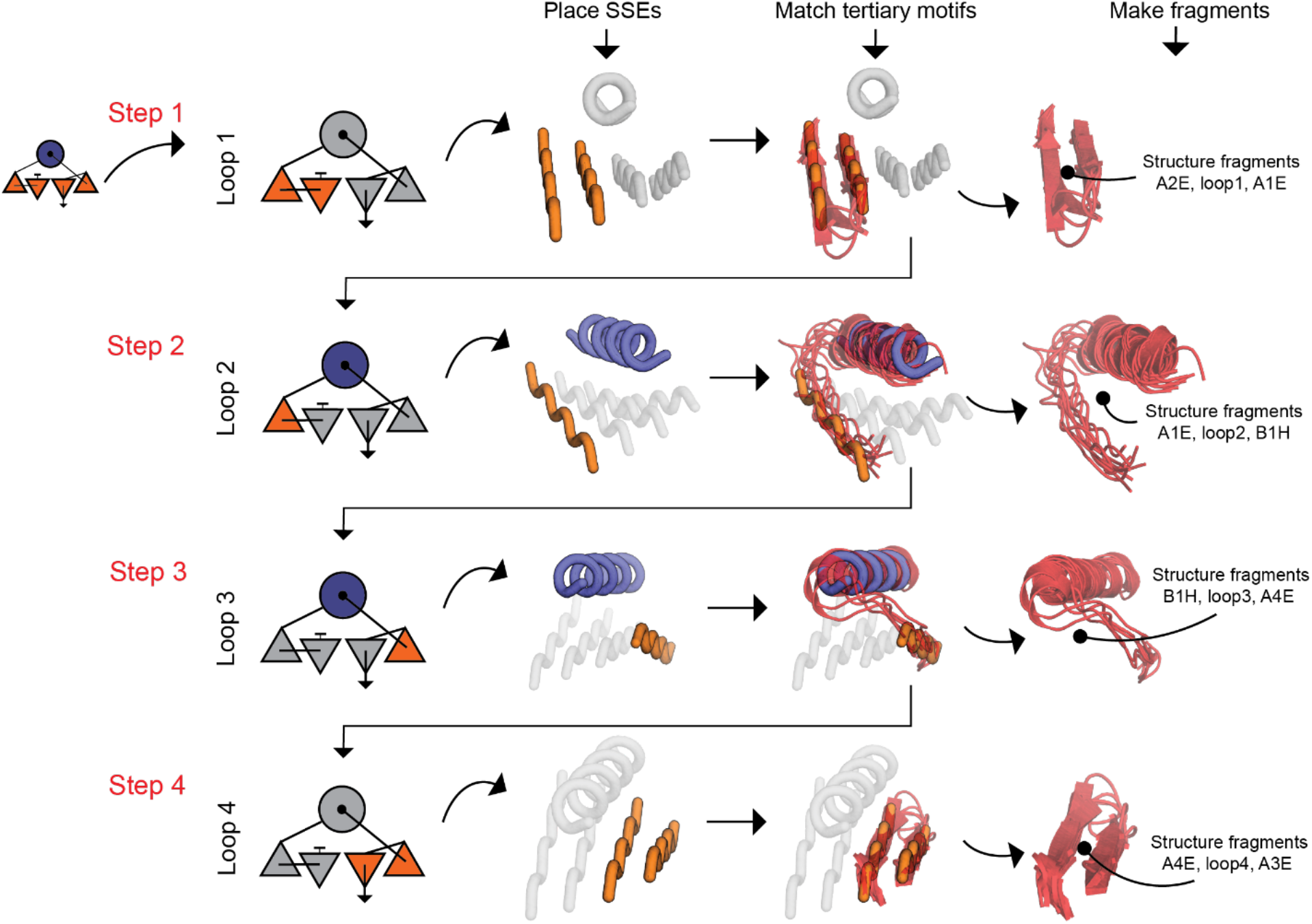
Loop assembling and fragment generation. Loop search and fragment generation for a three-layer fold. The loop assembler uses pairs of SSEs and searches for matching native tertiary motifs which are used to generate structural fragments of sizes 3 and 9 (3mers and 9mers).

**Supplementary Figure 5.**
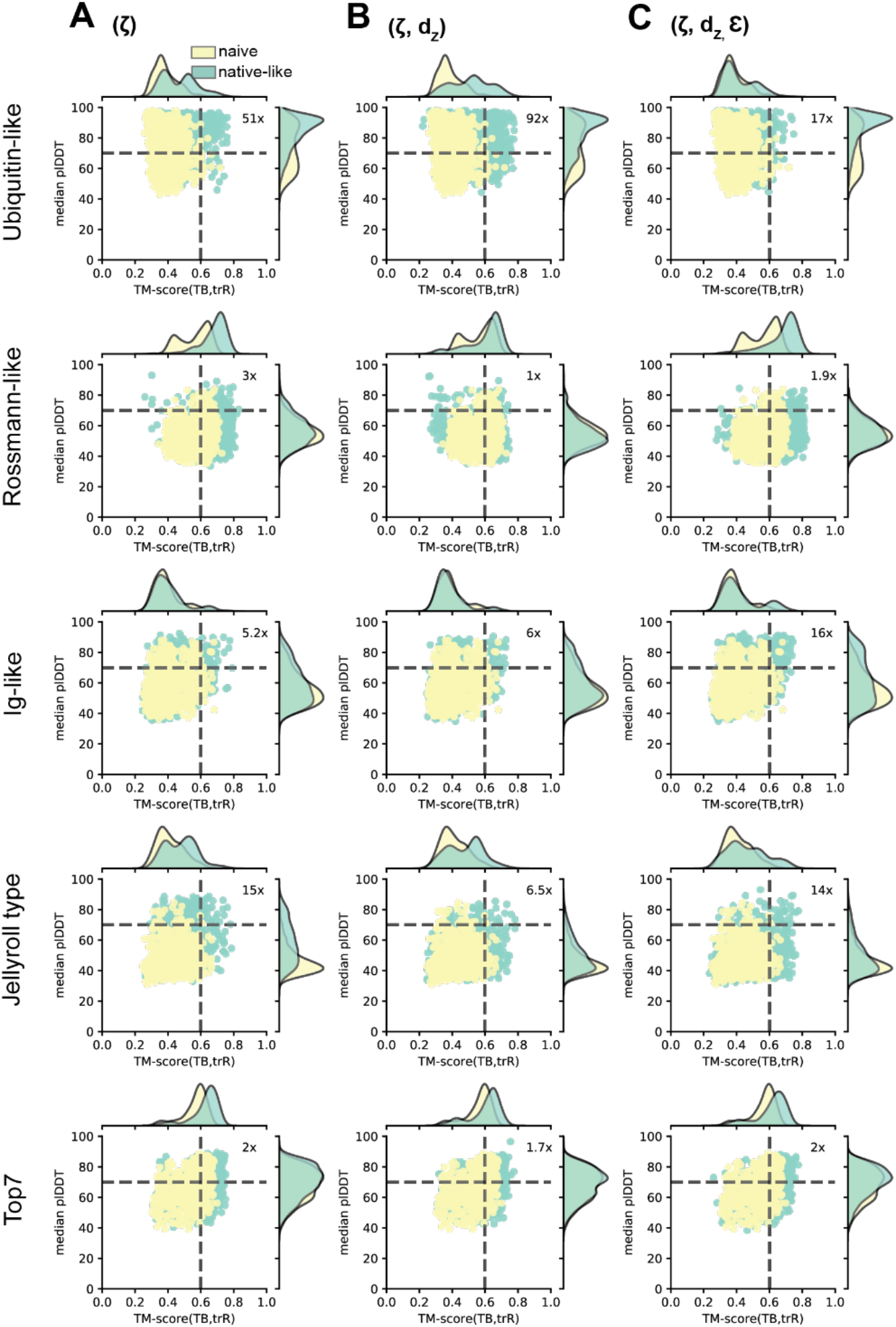
Influence of the different topological correction parameters on the quality of the designed sequences using TM-scores from trRosetta predictions and confidence scores from AlphaFold predictions. Sequence quality assessment by comparing the TM-scores calculated between the TopoBuilder (TB) models and the trRosetta (trR) models and the median AlphaFold (AF) predicted confidence score (median plDDT). **A:** Scenario using the twist angle ζ correction. **B:** Scenario using ζ + d_z_ corrections. **C:** Scenario with ζ + d_z_ + ε correction parameters.

**Supplementary Figure 6.**
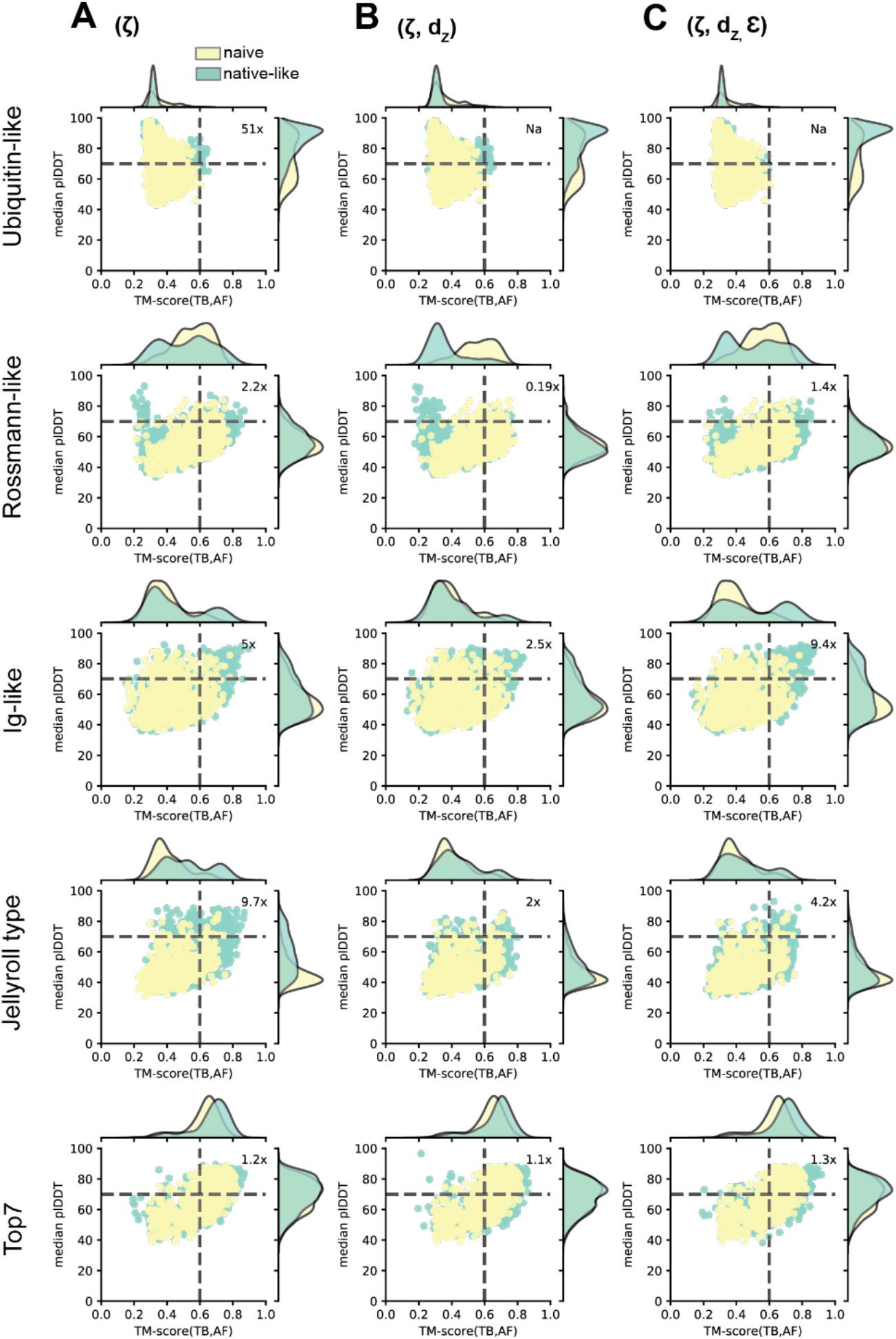
Influence of the different topological correction parameters on the quality of the designed sequences using TM- and confidence scores from AlphaFold predictions. Sequence quality assessment by comparing the TM-scores calculated between the TopoBuilder (TB) models and the AlphaFold (AF) models and the median AlphaFold predicted confidence score (median plDDT). **A:** Scenario using the twist angle ζ correction. **B:** Scenario using ζ + d_z_ corrections. **C:** Scenario with ζ + d_z_ + ε correction parameters.

**Supplementary Figure 7.**
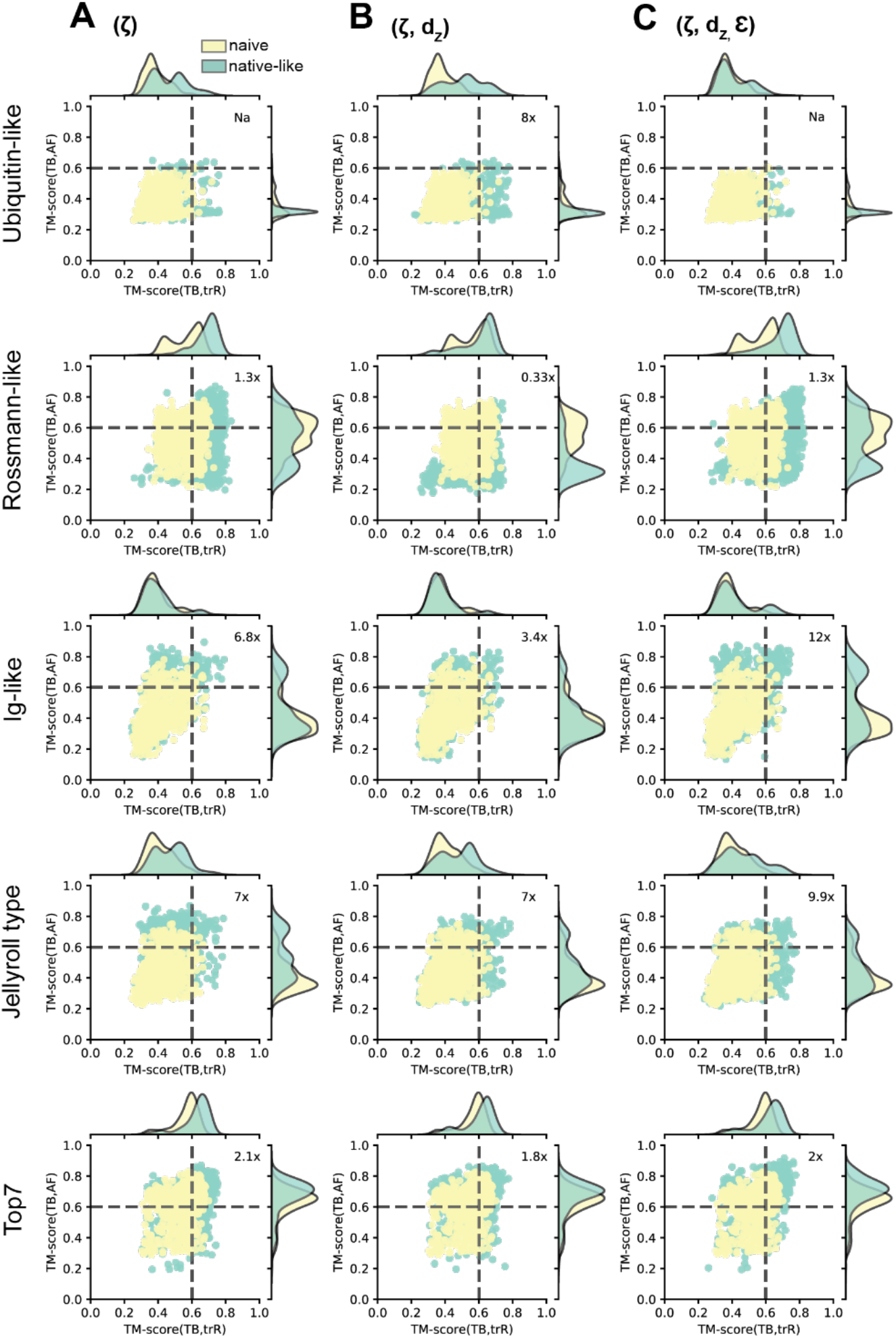
Influence of the different topological correction parameters on the quality of the designed sequences using TM- scores from AlphaFold and trRosetta predictions. Sequence quality assessment by comparing the TM-scores calculated between the TopoBuilder (TB) models and the AlphaFold (AF) models or trRosetta (trR) models. **A:** Scenario using the twist angle ζ correction. **B:** Scenario using ζ + d_z_ corrections. **C:** Scenario with ζ + d_z_ + ε correction parameters.

**Supplementary Figure S8.**
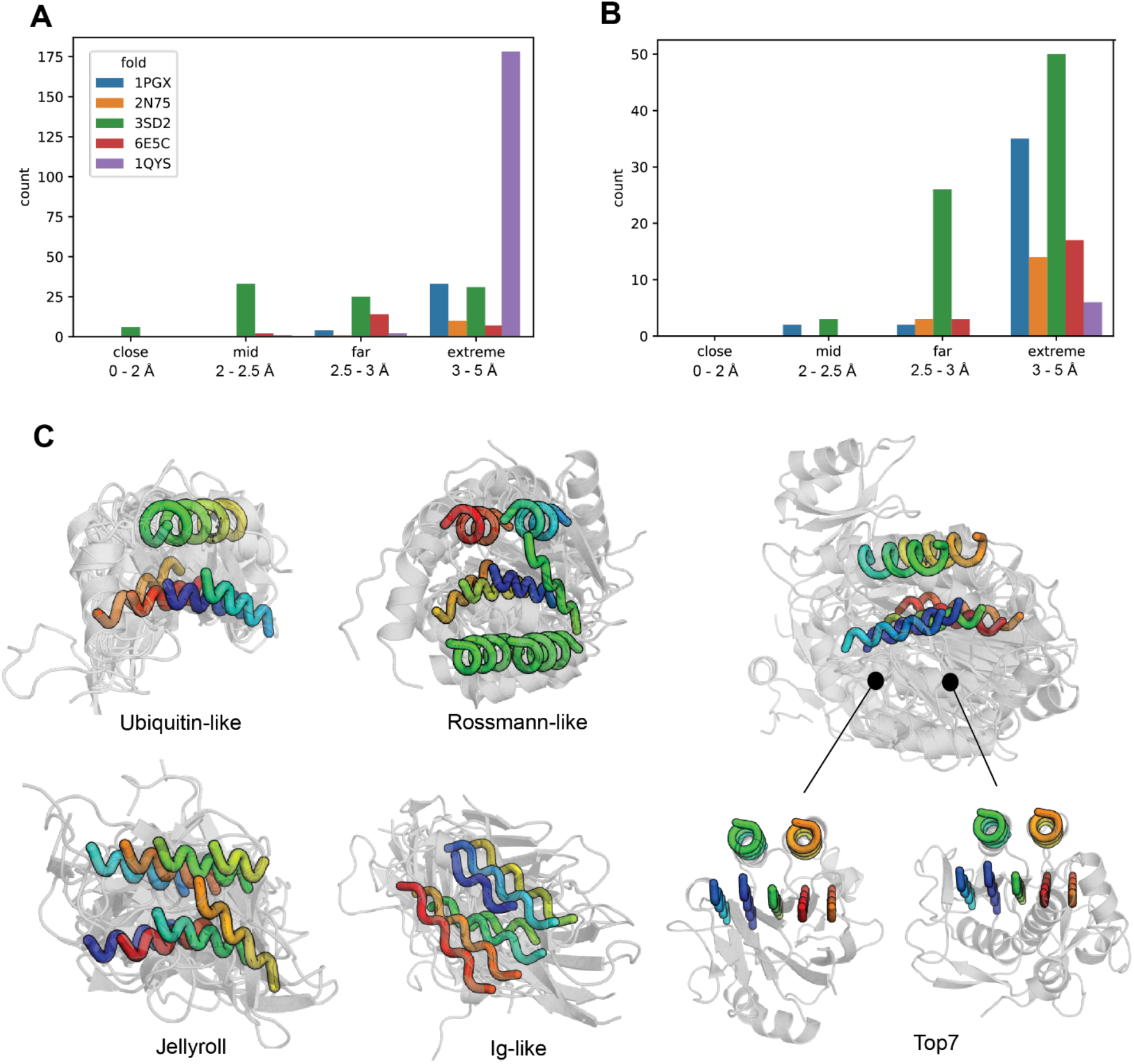
Counts and examples for the correction search. **A**: Counts of the retrieved matches per RMSD-bin (close: 0-2 Å, mid: 2-2.5 Å, far: 2.5-3 Å, extreme: 3-5 Å) for the first layer only (first step corrections). As only a single layer is searched more matches are retrieved. **B:** Counts of the retrieved matches per RMSD-bin for the second layer (second step corrections). Here, two layers are included making the matching more stringent, and less structures were retrieved. **C**: Examples of retrieved matches (second step corrections) to derive the geometrical corrections.

**Supplementary Figure S9.**
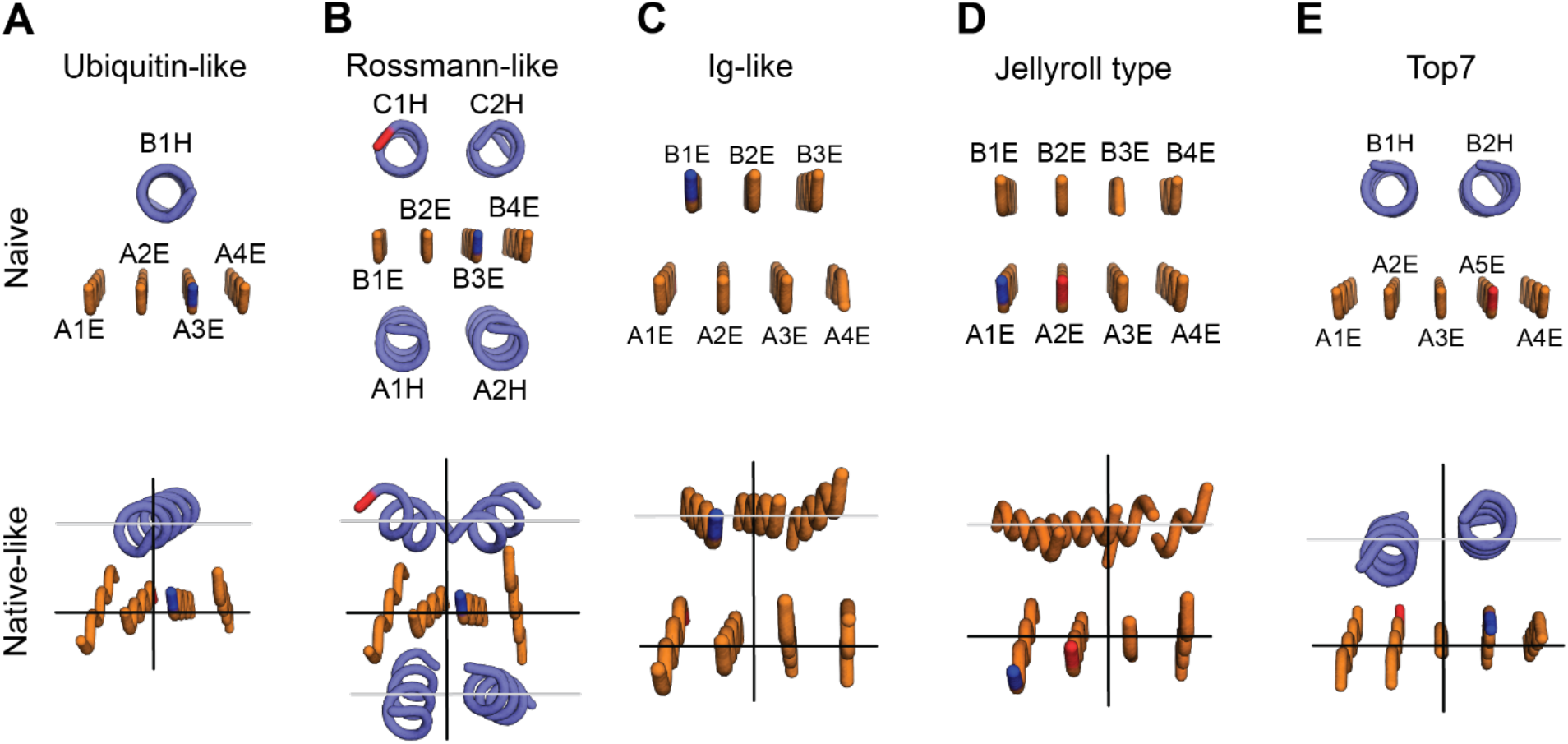
Comparison of naive and native-like Sketches. A front-view of naive and native-like Sketches. **A, B, C, D, E:** The naive Sketches (top row) with their respective Form element labeled for each of the 5 folds. The lower row shows the native-like Sketches with corrected SSE arrangements upon the TopoBuilder refinement.

**Supplementary Figure S10.**
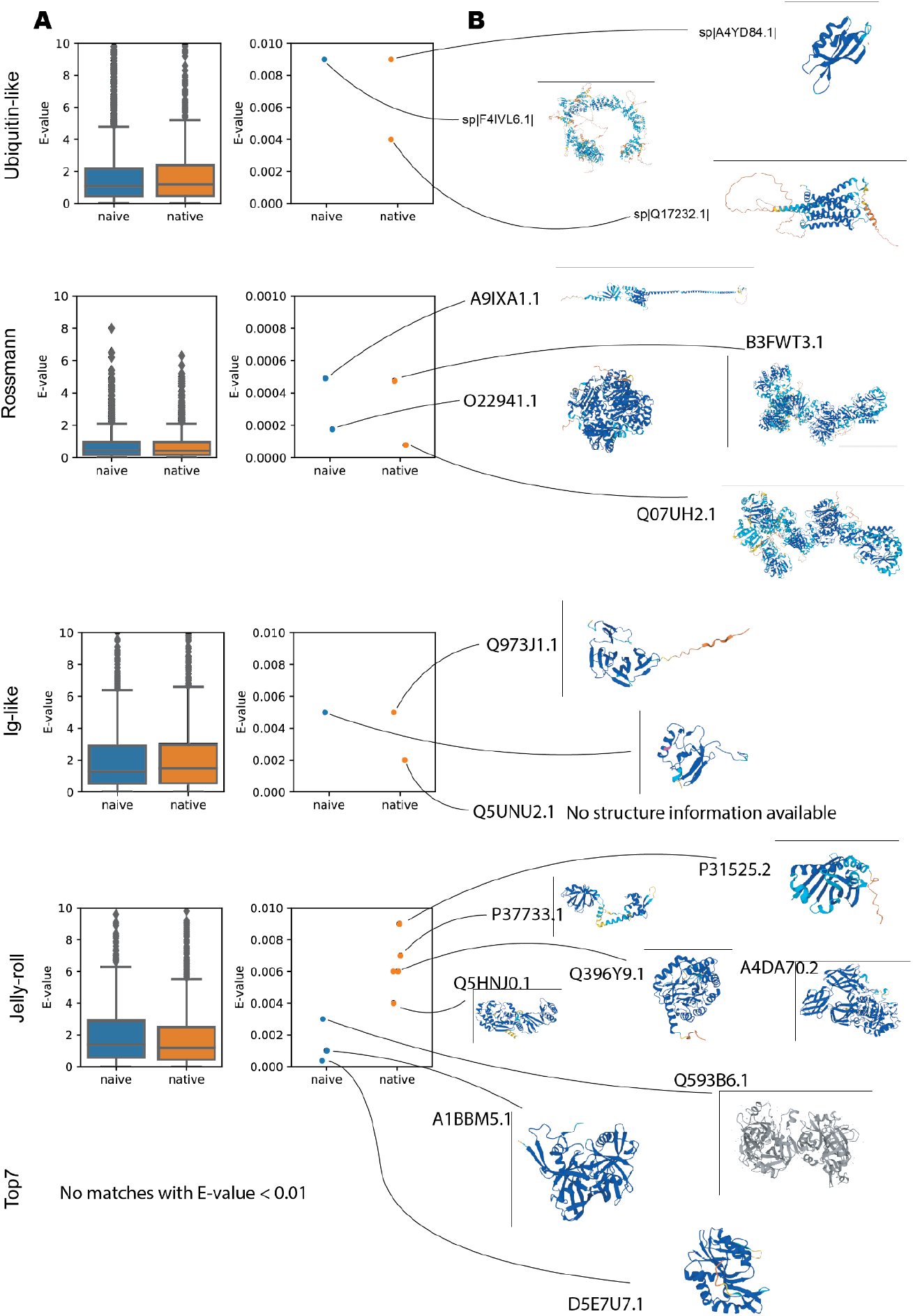
Blast search against SwissProt. **A:** Best E-values of the Blast searches against the SwissProt database for each design from a target fold. **B:** Available AlphaFold models (colored) or solved structures (grey). None of the models or structures presents a similar structure to the target fold.

**Supplementary Figure S11.**
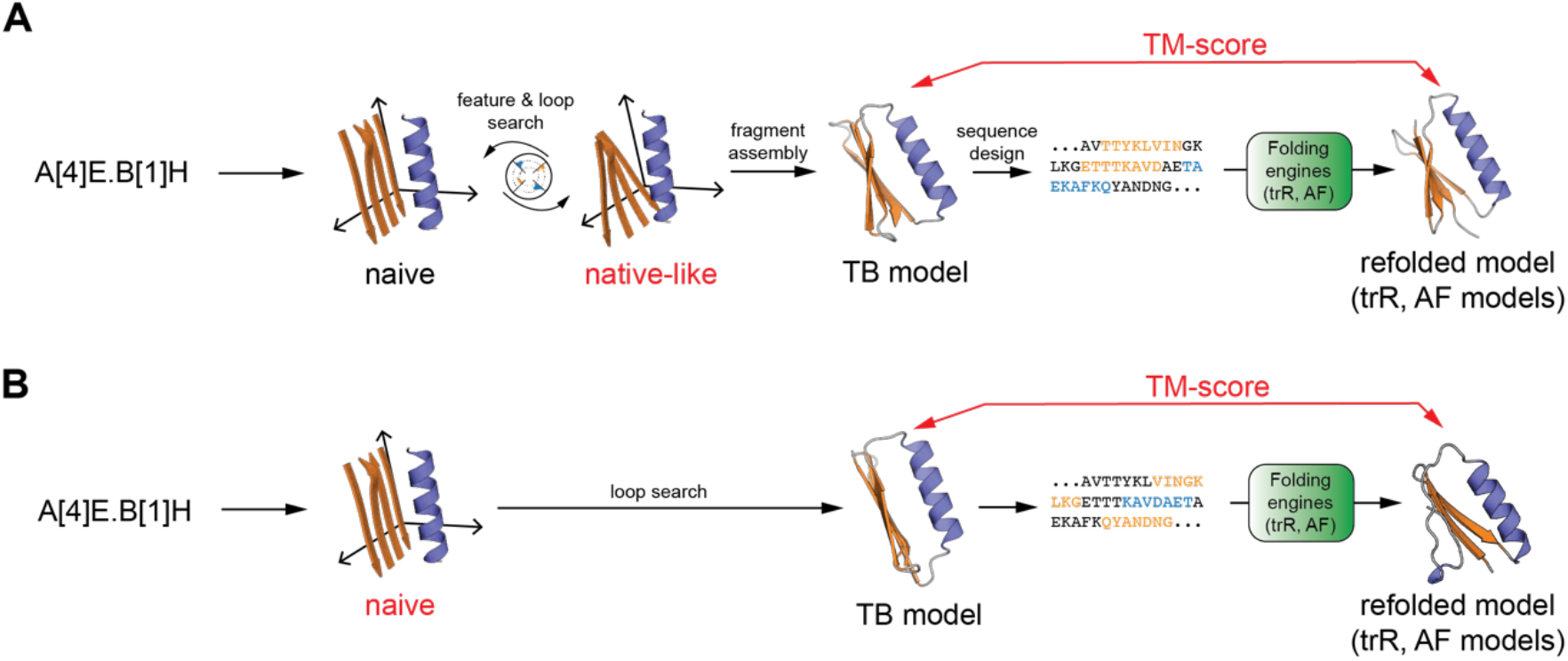
TopoBuilder *de novo* design pipeline. **A**: The TopoBuilder *de novo* design pipeline including topological correction step rendering a naive Sketch into a native-like structure with improved designability. A full atomistic model is generated through the Rosetta fragment assembly protocol (Rosetta FunFolDes) and sequences are designed using the Rosetta FastDesign method. To evaluate if our sequences encode the necessary fold determining signatures, we use state-of-the-art protein structure prediction engines (trRosetta and Alphafold). to computationally predict a model structure that we then structurally compare to our TopoBuilder (TB) model by calculating the TM-score. **B**: The TopoBuilder *de novo* design pipeline where the feature search module has been ablated to investigate the importance of the topological corrections.

**Supplementary Figure S12.**
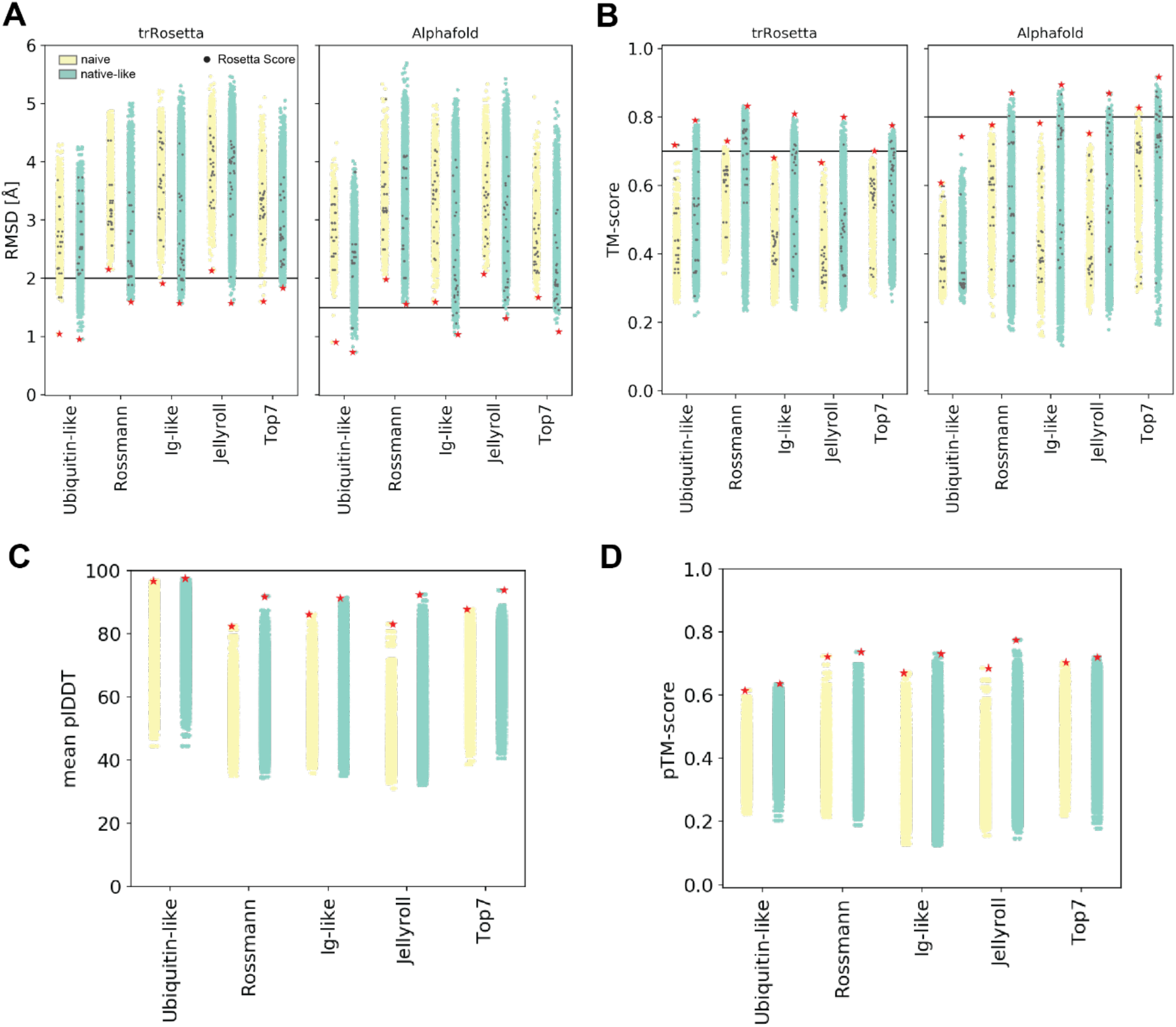
Assessment of the quality of the designed sequences using structure prediction tools. **A:** Pairwise TM-scores calculated between the TopoBuilder (TB) and the trRosetta or AlphaFold models (single sequence input without MSA) for each set of simulations. The black dots represent the 25 lowest scoring decoys by Rosetta energy. The red star indicates the best scoring decoy (either by TM-score or RMSD) for each of the simulations. **B:** RMSD using the best superposition and residue coverage between the TopoBuilder (TB) and the trRosetta or AlphaFold models. **C, D:** The mean plDDT and pTM-scores predicted by AlphaFold for each decoy.

**Supplementary Figure S13.**
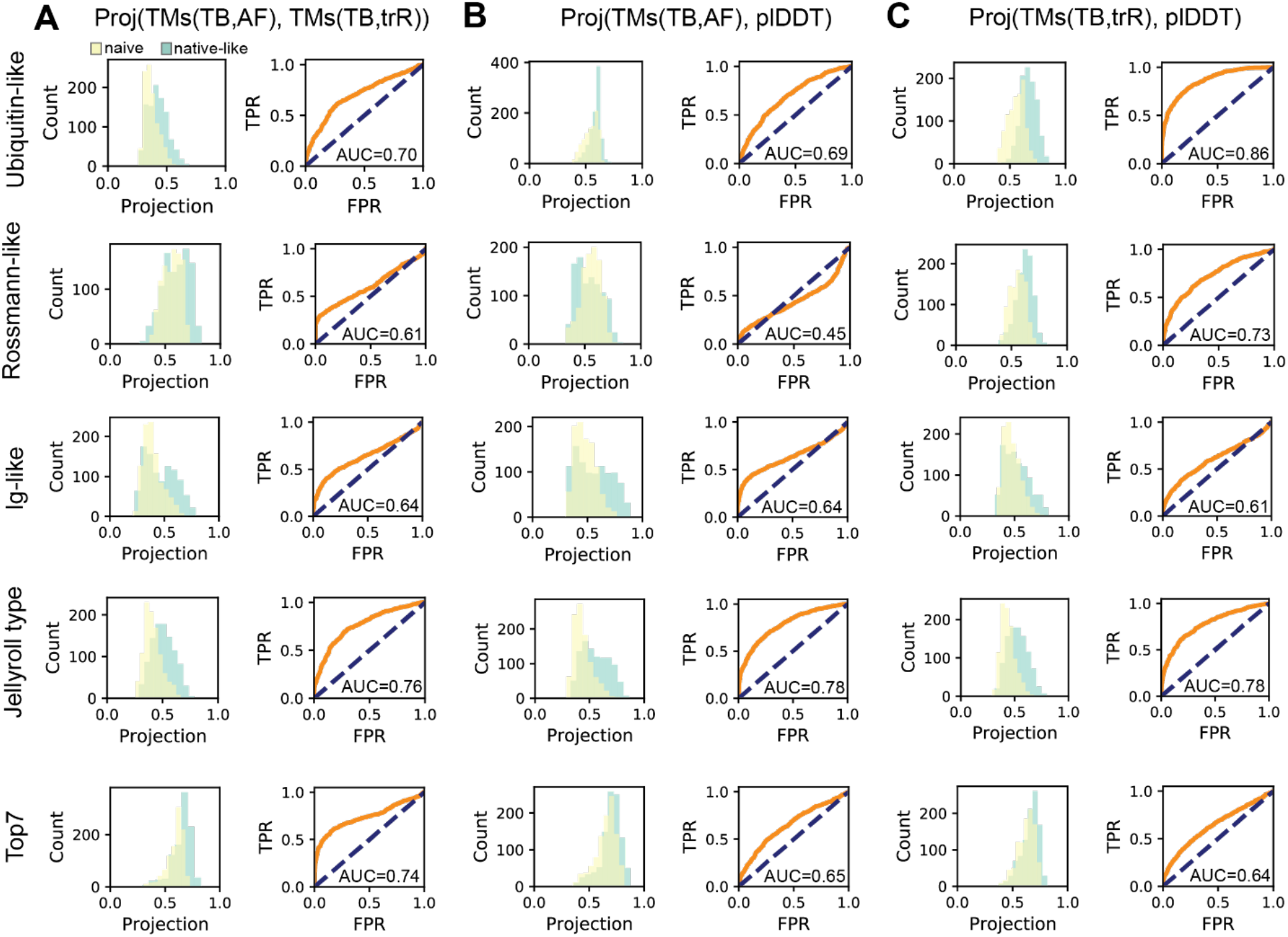
Sequence quality for various thresholds. The double-positive scores are projected onto the diagonal. The diagonal distributions are shown on the left, and the receiver operating characteristics (ROC) curve and the ROC area under the curve (AUC) are shown on the right evaluating the separation between the naive- and native-derived diagonal distributions. For example, if the AUC is 0.7, it means that there is a 70% higher chance of a native-derived sequence to lie towards the upper right quadrant than a naive-derived sequence. **A:** Projection onto diagonal of the {TM-score(TB,AF), TM-score(TB,trR)} scores. **B:** Projection onto diagonal of the {TM-score(TB,AF), plDDT} scores. **C:** Projection onto diagonal of the {TM-score(TB,trR), plDDT} scores.

**Supplementary Figure S14.**
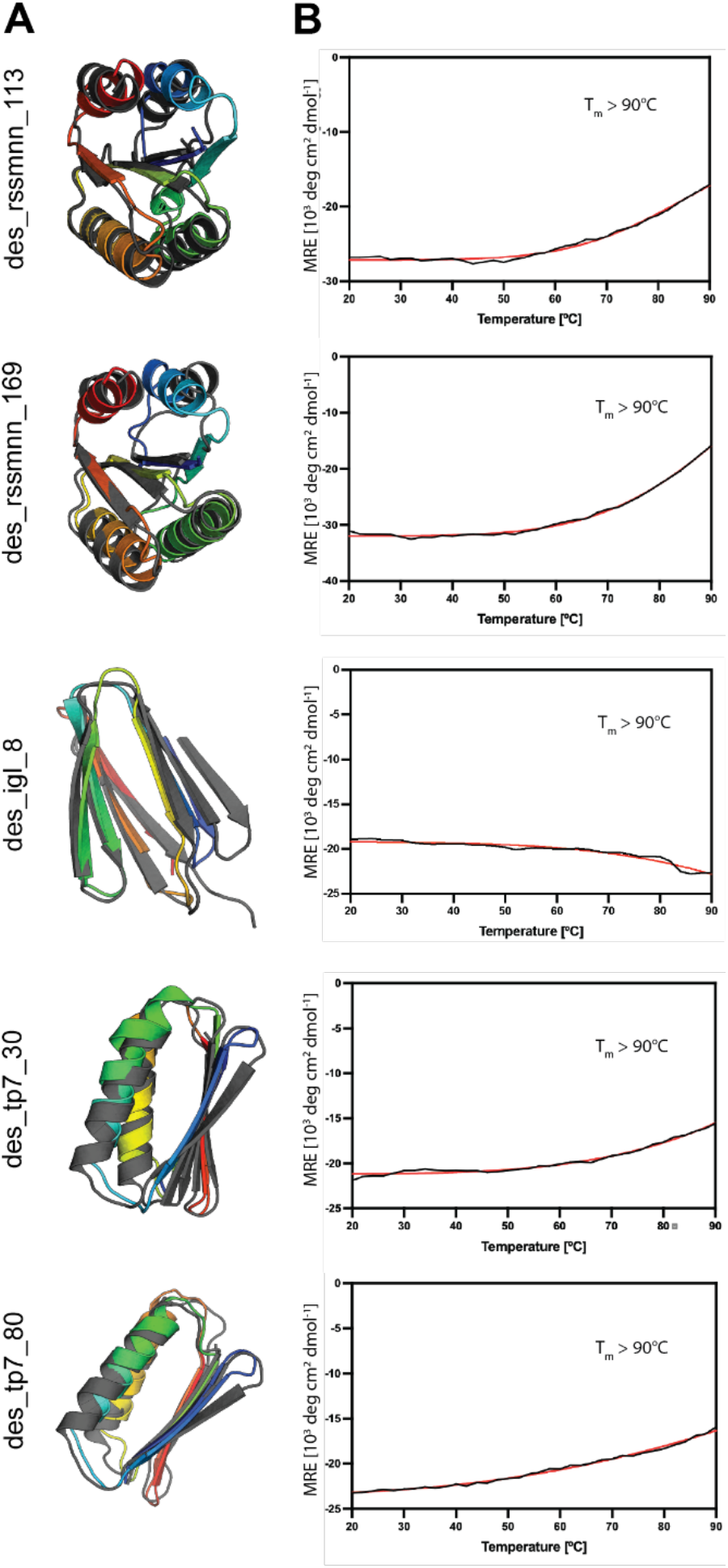
Thermal melting temperature determined by circular dichroism for the designed proteins. **A**: Models of the designs (rainbow) with their corresponding AlphaFold predictions (black). **B**: Thermal denaturation curves at wavelength of 220 nm for des_rssmnn_113, des_rssmnn_169, des_tp7_30 and des_tp7_80 and at 218 nm for des_igl_8 are shown.

**Supplementary Figure S15.**
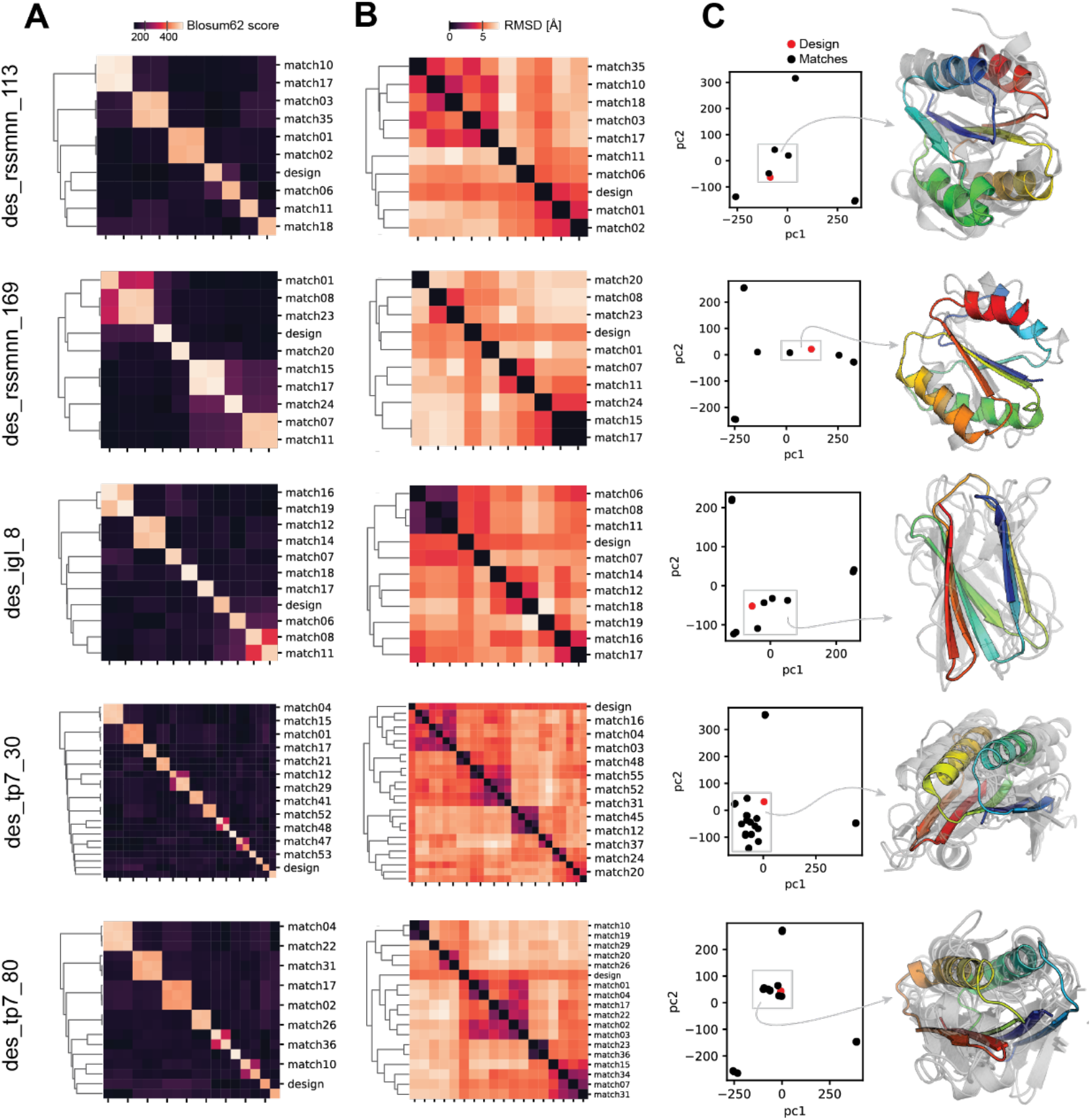
Sequence and structure similarities of the TB designs to native protein folds within the PDB. Comparison of the top MASTER matches to the TB designs on pairwise sequence, structure, and sequence + structure levels. **A**: Global pairwise sequence alignments scored with a Blosum62 distance (sum of the individual Blosum62 scores). **B**: Pairwise structure RMSDs. **C**: Principal component analysis (PCA) of the combined sequence (Blosum62 distance) and structure (RMSDs) features. Models of the designs (rainbow) with their cluster members.

**Supplementary Figure S16.**
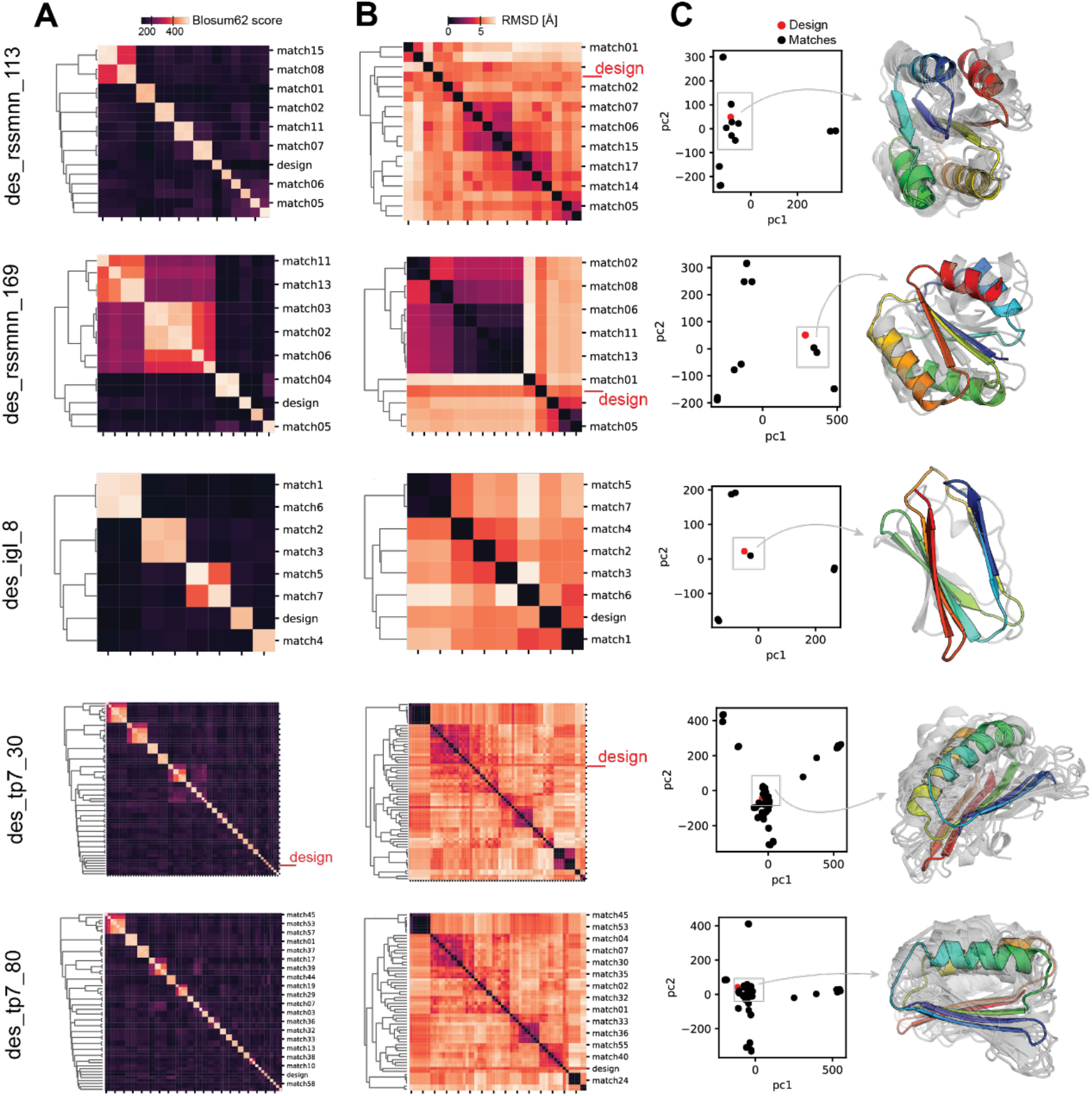
Sequence and structure similarities of the TB designs to the AF database. Comparison of the top MASTER matches to the TB designs on pairwise sequence, structure, and sequence + structure levels. **A**: Global pairwise sequence alignments scored with a Blosum62 distance (sum of the individual Blosum2 scores). **B**: Pairwise structure RMSDs. **C**: Principal component analysis (PCA) of the combined sequence (Blosum62 distance) and structure (RMSDs) features. Models of the designs (rainbow) with their cluster members.

## Supplementary tables

**Supplementary Table S1.**
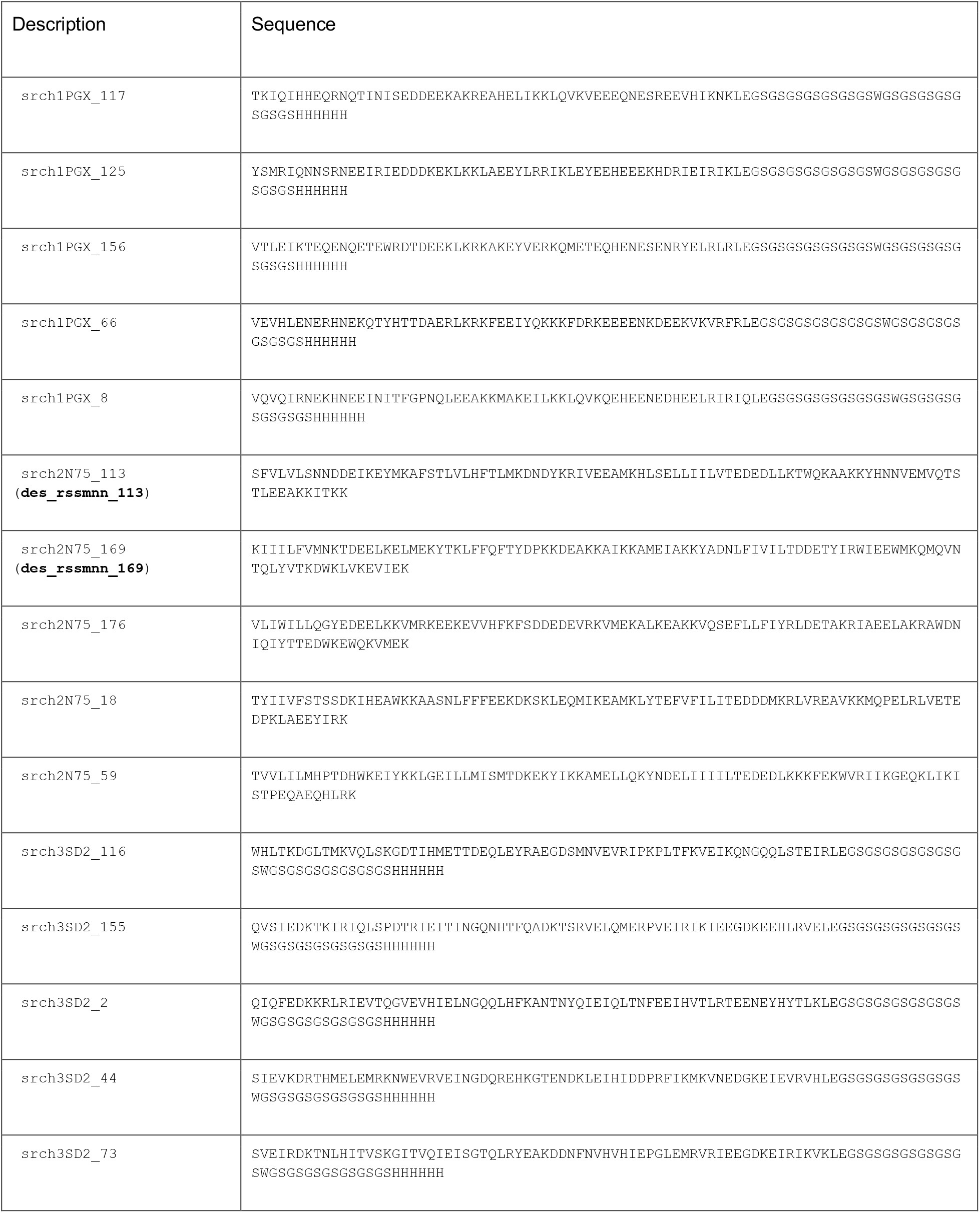
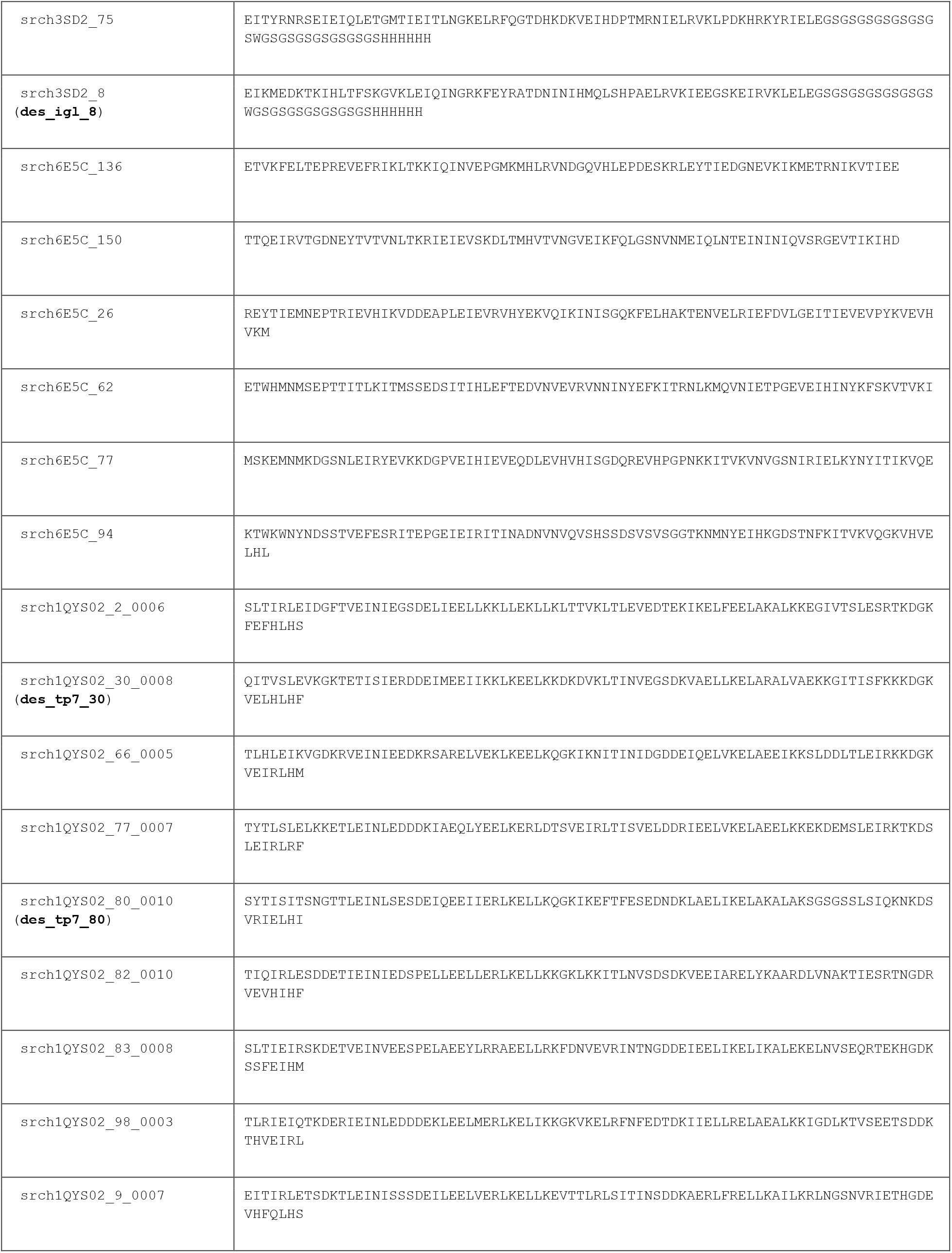
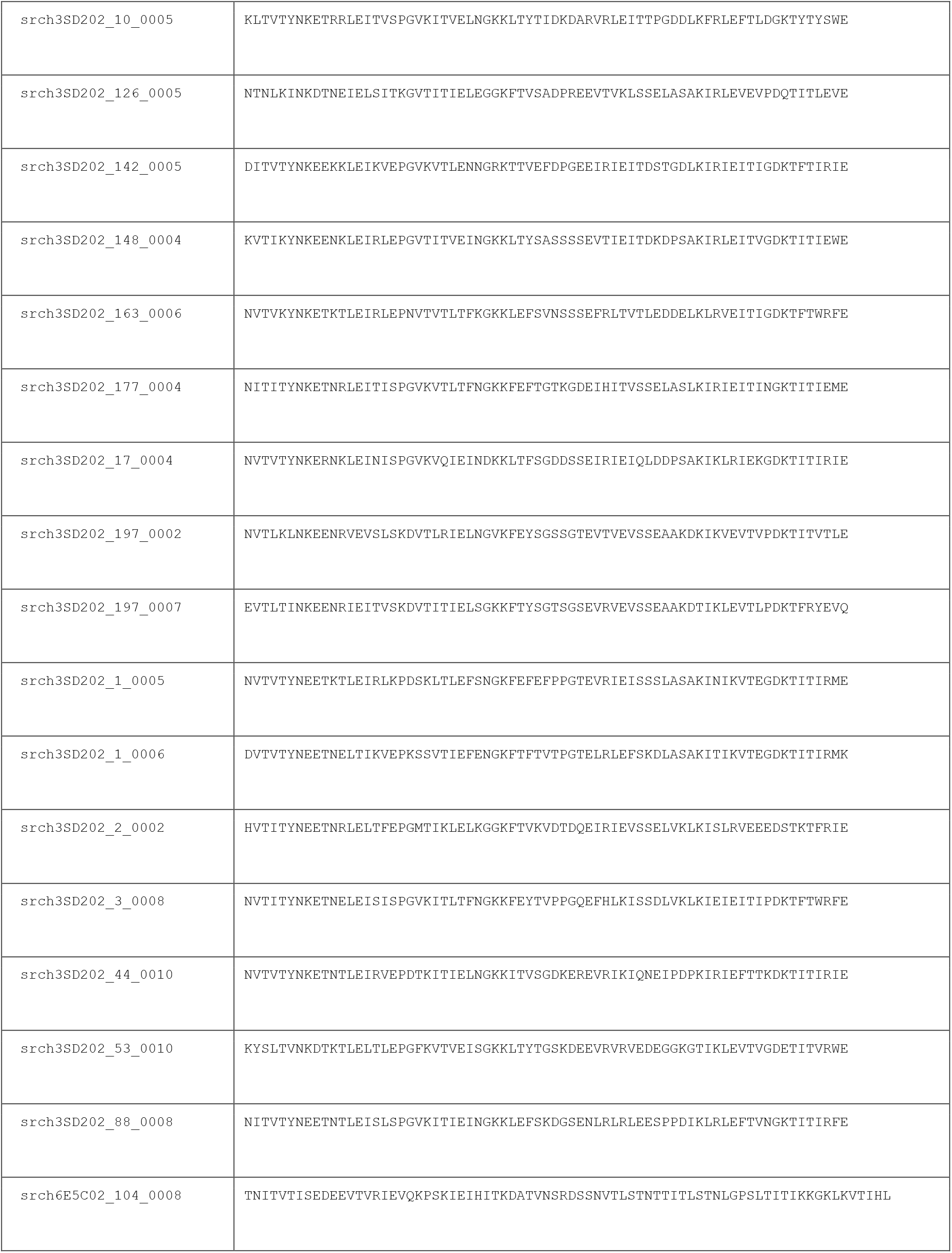
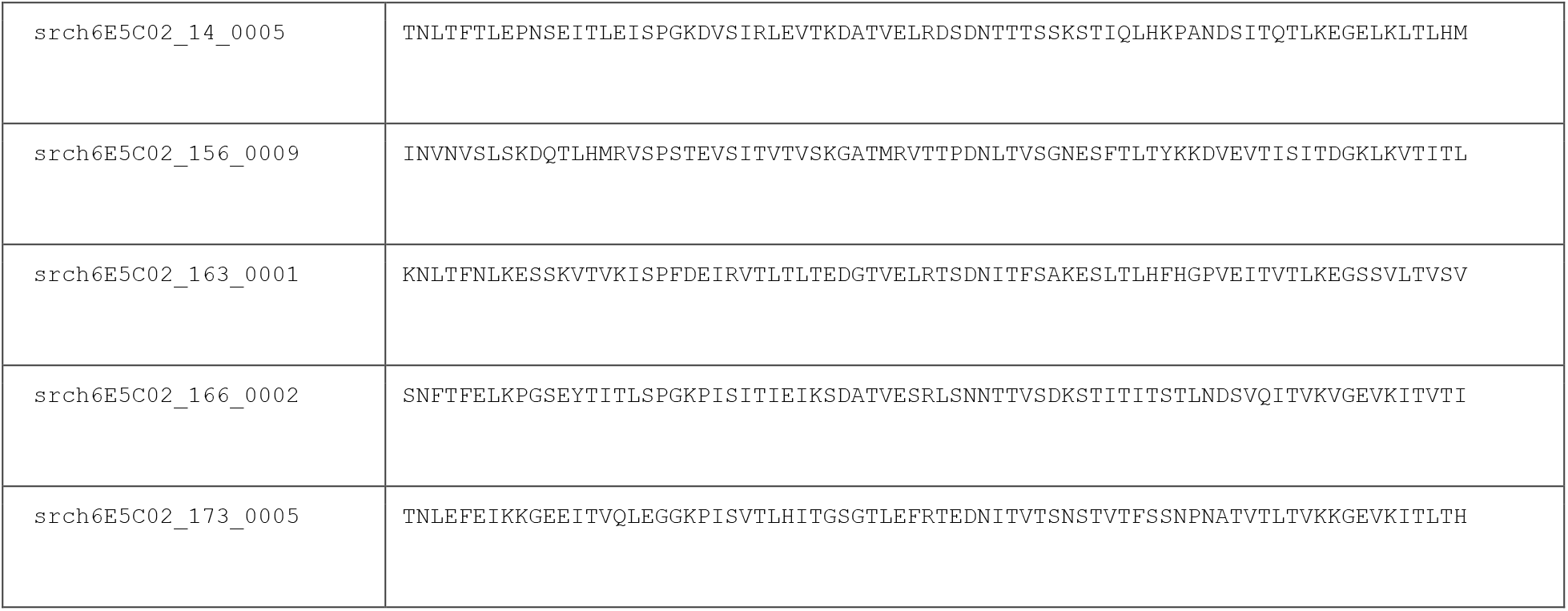
Designed sequences using the TopoBuilder.

